# State of India’s Birds 2023: A framework to leverage semi-structured citizen science for bird conservation

**DOI:** 10.1101/2024.09.05.611348

**Authors:** Ashwin Viswanathan, Karthik Thrikkadeeri, Pradeep Koulgi, J Praveen, Arpit Deomurari, Ashish Jha, Ashwin Warudkar, Kulbhushansingh Suryawanshi, MD Madhusudan, Monica Kaushik, Naman Goyal, Priti Bangal, Rajah Jayapal, Suhel Quader, Sutirtha Dutta, Tarun Menon, Vivek Ramachandran

## Abstract

Birds and their habitats are threatened with extinction around the world. Regional assessments of the ‘State of Birds’ are a vital means to prioritize data-driven conservation action by informing national and global policy. Such evaluations have traditionally relied on data derived from extensive, long-term, systematic surveys that require significant resources, limiting their feasibility to a few regions in the world. In the absence of such ‘structured’ long-term datasets, ‘semi-structured’ datasets have recently emerged as a promising alternative in other regions around the world. Semi-structured data are generated and uploaded by birdwatchers to citizen science platforms like eBird. Such data contain inherent biases because birdwatchers are not required to adhere to a fixed protocol. An evaluation of the status of birds from semi-structured data is therefore a difficult task that requires careful curation of data and the use of robust statistical methods to reduce errors and biases. In this paper, we present a methodology that was developed for this purpose, and was applied to produce the comprehensive State of India’s Birds (SoIB) 2023 report. SoIB 2023 assessed the status of 942 bird species in India by evaluating each species based on three metrics: 1) long-term change: change in abundance between the year 2022 and the year-interval pre-2000; 2) current annual trend: mean annual change in abundance from 2015 to 2022; and 3) distribution range size. We found evidence that 204 species have declined in the long term, and 142 species are currently declining. We present and discuss important insights about India’s birds that can guide research and conservation action in the region. We hope that the detailed methodology described here can act as a blueprint to produce State of Birds assessments from semi-structured citizen science datasets and springboard conservation action in many other regions where structured data is lacking but strong communities of birders exist.

**Open Research Statement:** The primary data are already publicly available on eBird (Sullivan et al. 2014), and other data are already published in a GitHub repository (stateofindiasbirds 2024).

## Introduction

Birds are rapidly declining worldwide (BirdLife International 2022b, Lees et al. 2022), mirroring a broader decline in global biodiversity (Díaz et al. 2019). Reasons for the decline, and potential solutions, vary with species and geopolitical region (BirdLife International 2018). As opposed to global evaluations, regional assessments of bird population status are often better placed to effectively guide conservation and species recovery action (BirdLife International 2015, 2017, Díaz et al. 2019, BirdLife International 2022b). Bird population status assessments (State of Birds reports) have been published in the past decade for several countries and regions (e.g., Turkey, Kittelberger et al. (2023); North America, North American Bird Conservation Initiative (2022); India, SoIB (2020); United Kingdom, Hayhow et al. (2017); Switzerland, Moosmann et al. (2023); Australia, BirdLife Australia (2015b); and Europe, BirdLife International (2015)), helping guide national conservation priorities and policy.

Systematic surveys form the backbone of typical ‘State of Birds’ reports. Such surveys are usually conducted periodically (often every year) by trained volunteers who use fixed protocols and collect ‘structured’ data. They follow a sampling design that ensures similar sampling effort across space (across the country/region) and time (over years), allowing straightforward inference about bird population trends across large regions. Some examples of such surveys include the New Atlas of Australian Birds (Barrett et al. 2003, BirdLife Australia 2015a), United Kingdom Breeding Bird Survey (Harris et al. 2018, British Trust for Ornithology 2019), North American Breeding Bird Survey (Sauer et al. 2013), Pan European Common Bird Monitoring Scheme (European Bird Census Council (EBCC) 2018) and Swiss Common Breeding Bird Survey (Knaus et al. 2018). These systematic surveys at the scales of entire countries or regions, however, are resource- and training-intensive, and are currently infeasible in many countries in the Global South where requisite resources and volunteer networks may be lacking.

A new opportunity for regional assessments has now emerged in such countries. This comes from data collected by hobbyist birdwatchers, who go out and observe birds for pleasure.

Birdwatchers have a long-standing tradition of recording checklists of birds they observe. Previously, they would record these lists in notebooks but today, the lists can be directly uploaded to citizen science platforms like eBird (Sullivan et al. 2014), creating vast repositories of information that are publicly accessible. Data uploaded by birdwatchers to platforms like eBird, however, are ‘semi-structured’ because they are generated from birdwatching sessions. Birdwatchers are allowed some flexibility in protocol when making lists, introducing biases into observation effort (time spent birdwatching, distance travelled, etc.) and heterogeneity in space and time. These biases complicate the process of curating, analysing, and inferring large-scale abundance trends from such data.

State of Birds assessments typically use species count data to derive periodic estimates of population size (e.g., Gregory et al. 2005, Newson et al. 2005, Herrando et al. 2008, Newson et al. 2008, Blancher et al. 2009, Brotons and Herrando 2011, Norman et al. 2012, Musgrove et al. 2013, Crowe et al. 2014, North American Bird Conservation Initiative 2014, Sattler et al. 2017). Species counts, however, may not be reliable in semi-structured data generated from recreational birdwatching because counting methods and attention to detail tend to vary between observers. An alternative to species counts in semi-structured data is species detection/non-detection data. Detection/non-detection data can be used to derive periodic estimates of a species’ ‘reporting frequency’, i.e., the average likelihood of encountering a species given a specified effort.

Periodic assessments of both population size and reporting frequency can be used to estimate abundance trends over time. Both methods produce trends that are often correlated (Joseph et al. 2006, Pollock 2006), and detection/non-detection data have already been employed to estimate trends for State of Birds assessments (Olsen et al. 2003, Szabo et al. 2010, BirdLife Australia 2015b, a.)

With the rapid growth in popularity of eBird in India since 2014 and the emergence of a vast dataset, it recently became feasible to assess Indian bird species trends using detection/non-detection data. A partnership of organizations used data uploaded to eBird to produce the State of India’s Birds 2020 report (SoIB 2020). Since the publication of the 2020 report, the eBird database has rapidly grown in size and quality, allowing the partnership to produce the next, and more robust, status assessment of birds in the country, State of India’s Birds 2023 (SoIB 2023a, b.). Here, we describe the key findings of SoIB 2023, and the detailed methods used to derive its various components from semi-structured data uploaded to eBird. We describe in detail the steps used to filter data to reduce inherent biases, and the rationale used to select species for inclusion in various analyses. We also highlight analytical steps used to reduce spatial autocorrelation and to account for variation in birdwatching effort between checklists. The analytical framework described here can be applied to produce a State of Birds assessment in any other region where long-term semi-structured data is available.

## Methods

### Introduction to eBird data

Birdwatchers upload their observations to eBird through the basic unit of a checklist. A checklist can contain zero to several species of birds listed alongside their counts, where every species is named according to the Clements taxonomy (Clements et al. 2023). A checklist is traditionally uploaded using one of four protocols, ‘incidental’, ‘stationary’, ‘traveling’, or ‘historical’. The incidental protocol is used for a list of birds observed when not primarily watching birds; only date and location inputs are mandatory. The other three—stationary, traveling and historical— are used for checklists generated during time devoted to birdwatching. Apart from date and location, birdwatching effort must be specified for stationary (start time and duration) and traveling checklists (start time, duration, and distance). The historical protocol is used when this information on effort is not available, which is often the case with data collected years ago, and for example, transcribed from physical notes. More information on the protocol types is on the eBird help pages (eBird 2024). When uploading a checklist, birdwatchers are asked whether the list is ‘complete’. For a checklist to be marked complete, it should include every species identified by the birdwatcher (seen or heard) in the duration of the checklist. This is essential information because only complete checklists allow an analyst to infer non-detection, and therefore only these checklists can be used to calculate statistically sound reporting frequencies and trends.

### Components of SoIB 2023

State of Birds (and similar) reports usually include classifications of bird species into colour-coded concern or priority categories based on their population trends (North American Bird Conservation Initiative 2012, BirdLife Australia 2015b, a, Eaton et al. 2015a, b, Hayhow et al. 2017). Such categorization enables prioritization of species for research and conservation action. We developed a methodology to assess the Conservation Priority status of a species (High, Moderate, Low) based on three metrics estimated from eBird data: 1) long-term change (LTC): change in abundance between the year 2022 and the year-interval pre-2000; 2) current annual trend (CAT): mean annual change in abundance from 2015 to 2022; and 3) distribution range size (DRS). These three metrics and the Conservation Priority status derived from them form the backbone of SoIB.

### The SoIB pipeline

The SoIB pipeline, or the analytical workflow used to produce the essential output of SoIB, is outlined stepwise in **Fig. 1**. The pipeline includes several steps from downloading and curating the data to calculating the three metrics that are used to derive Conservation Priority status. The statistical software R 4.2.3 (R Core Team 2023) was used for all steps in the pipeline.

**Fig. 1:**
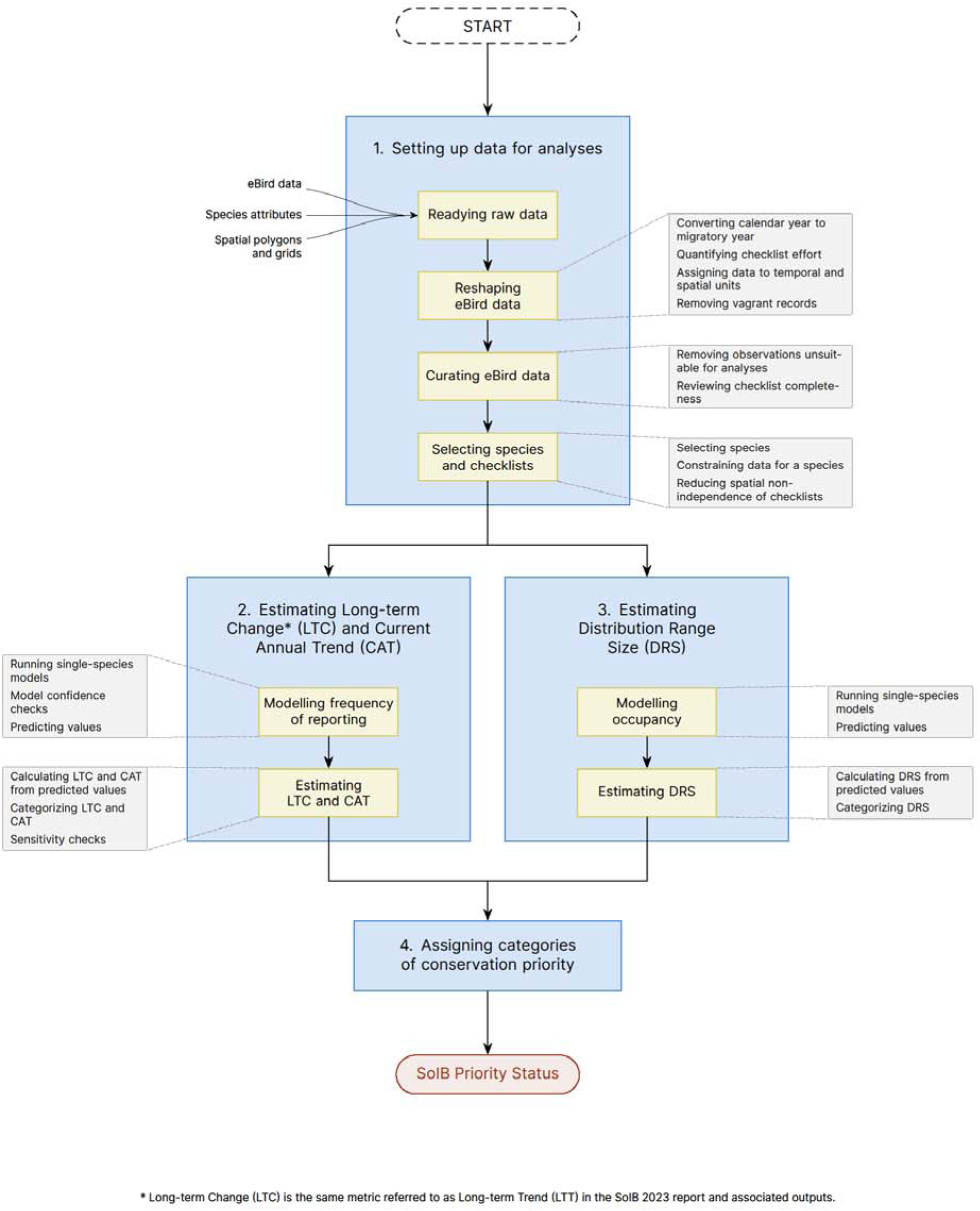
The SoIB pipeline. Original design by Janhavi Rajan.

#### Setting up data for analyses

Data downloaded from eBird require additional curation to be ready for analyses. This section describes how eBird data was reshaped and curated, after appending species-level and spatial polygon information.

##### Readying raw data

SoIB 2023 used three kinds of data: eBird data; species-level information including habitat specialization, endemic region, etc.; and spatial data in the form of administrative and gridded maps of India.

##### eBird data

As the basis for the report, we used the ‘relMay-2023’ version of the eBird Basic Dataset (EBD) for India that contains all data submitted up to 31 May 2023 (downloaded on 15 June 2023, follows taxonomy in Clements et al. 2022). We separately requested eBird for ‘sensitive’ species data which is not included in the public downloads. The EBD and sensitive species data together made up the base dataset required for our analyses (**Fig. 1**).

##### Species attributes

For the second kind of data, we used available knowledge from handbooks, field guides (Ali and Ripley 1968, Grimmett et al. 2011, Rasmussen and Anderton 2012), published and grey literature, and online resources to collate species-specific information for all of India’s bird species (1,358 as of 31 May 2023 on eBird). This information includes taxonomic groupings (Clements et al. 2022), IUCN status (IUCN 2023), diel activity classification (Wilman et al. 2014), diet guild (Wilman et al. 2014), endemism and region of endemism (adapted from Praveen and Jayapal 2023), and habitat specialization and migratory status within India (adapted from Wilman et al. 2014)—all also fine-tuned using other sources. We identified species that needed to be included in SoIB from the Indian perspective (birds for which India constitutes a large and/or important part of their breeding or wintering ranges, culturally significant birds, etc.), even if it did not meet the minimum data-driven criteria for species selection. Finally, we added information about whether a species’ range within India is restricted to islands (even if not an endemic). We used this set of information to filter data, select species, and aggregate species for analyses based on their taxonomic and ecological attributes (**Fig. 1**, 1. Selecting species and checklists - Selecting species). The sheet containing this species-level information can be downloaded from GitHub (stateofindiasbirds 2024).

##### Spatial polygons and grids

For spatial data, we used a shapefile of India’s administrative boundary to produce clipped (to India’s boundary) and unclipped square spatial grids of the following edge resolutions: 5 km, 25 km, 50 km, 100 km, 200 km. Since spatial gridding functions use fixed-degree cell size specifications, and due to the curvature of the earth that results in a variation in the degree:km relationship of longitude with latitude, the horizontal edges of our grid cells are shorter in northern latitudes than in southern latitudes. As a result, the longitudinal edges of our 25 km resolution, for instance, range from 24.8 km at the southernmost tip to 20 km at the northernmost tip. We also created a spatial polygon that included a 1 km buffer around the country’s boundary to be able to isolate data located in the ocean from data located inland and in the intertidal zones. An elevation raster was created for use as the background in species distribution range maps.

##### Reshaping eBird data

This section describes how the raw data at this stage were modified to make them suitable for curation and analyses. As a first step, we removed duplicate instances when a species name (observation) was associated with multiple shared checklists between observers from the same birdwatching party.

##### Converting calendar year to migratory year

We obtained information about the day, month, calendar year, and what we call the ‘migratory year’ of every observation, which is similar in concept to a ‘water year’ (Wikipedia contributors 2024). Winter migrants to the Indian subcontinent arrive in autumn and leave the next spring; a migratory cycle consequently spans two calendar years. We therefore shifted every calendar year forward by five months to create a migratory year, which encloses a single migratory cycle. Any given migratory year thus consists of the period between 1 June of one calendar year and 31 May of the subsequent (i.e., 2022 refers to the period 1 June 2022–31 May 2023). The word ‘year’ in this paper hereafter and in SoIB (2023b) refers to the migratory year so defined.

##### Quantifying checklist effort

Birdwatching effort can vary considerably between eBird checklists. Some checklists may involve just a few minutes of birding and others may involve several hours. The probability of reporting a bird in a checklist will depend on the effort spent birding - the greater the effort, the greater the probability. The reporting frequency of a species during a specific year-interval will therefore depend on the effort spent birding in all the checklists from that year. Because this effort does not remain consistent from year-interval to year-interval on eBird, it is necessary to standardize birdwatching effort between checklists so that there is no bias while making comparisons between years.

Birdwatching effort is traditionally specified in an eBird checklist by the maker of the checklist in two ways - by noting the duration and/or the distance travelled. Another measure called ‘list length’ has also sometimes been used as a proxy for birdwatching effort on eBird (Horns et al. 2018, Neate-Clegg et al. 2020). List length is the number of unique species reported across a set of shared checklists. Unlike duration and distance travelled, list length is not specified by the maker of a checklist. In this study, we used list length as a measure of birdwatching effort to statistically model frequency of reporting (**Fig. 1**, 2. Modelling frequency of reporting - Running single-species models) instead of duration and distance travelled. We arrived at this decision after examining the relationships between frequency of reporting and these three candidate explanatory variables for a number of species. We found, for 50 randomly selected species, that the frequency of reporting correlates poorly with duration or distance, indicating that they are unreliable predictors of reporting frequency (**Fig. 2, Fig S2-7**). The correlation with list length, however, is strong.

**Fig. 2:**
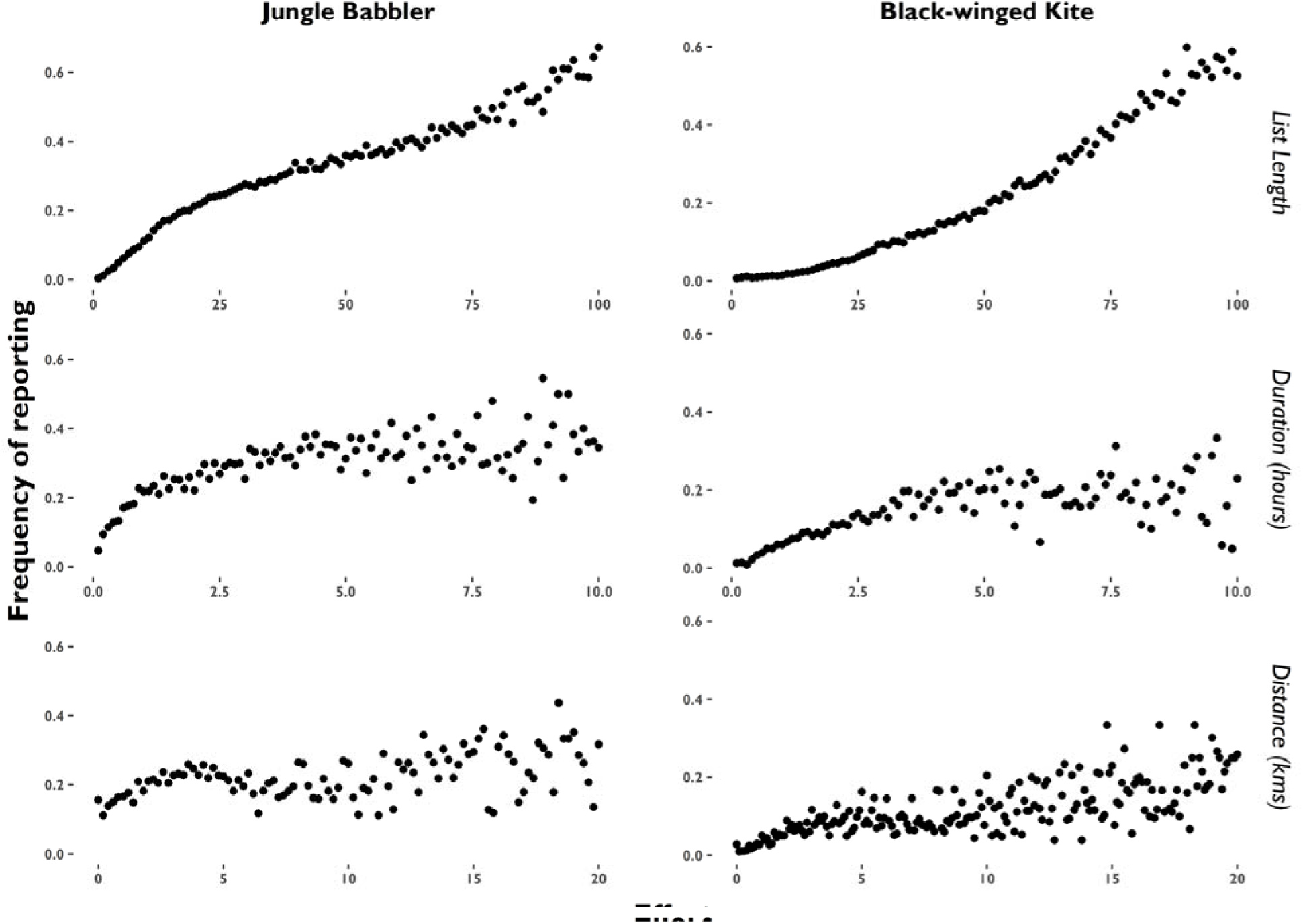
Frequency of reporting of (representative) Jungle Babbler and Black-winged Kite in relation to list length (number of species in a checklist up to 100), binned Duration (100 0.1 hrs bins), and binned Distance (100 0.2 km bins. Patterns were similar for 50 randomly selected species (**Fig S1-6**).

Duration and distance may be unreliable because they depend on individual birdwatching styles and individual comprehension of eBird best practices. For example, many birdwatchers document double the distance when they go up and down a single trail but others (correctly) report only the one-way distance. Similarly, a birdwatching list that spans 8 hours may actually only involve 2–3 hours of birdwatching, with the rest being time spent driving or, say, at food stops. The number of species on a checklist, however, correlates well with the effort put in, in spite of the obvious correlations also with bird diversity and detectability in a region (see section in supplement called “Relationship between reporting frequency and list length” and **Fig. S7**). Other recent studies have therefore used list length to explain birdwatching effort (Horns et al. 2018, Neate-Clegg et al. 2020). An additional critical advantage of using list length as a proxy of effort is that it allows the use of checklists from before 2010 uploaded with the historical protocol, for which duration and distance metadata are usually missing (**Fig. S1**).

##### Assigning data to temporal and spatial units

We organized the data into 14 year-intervals for LTC and CAT analysis: pre-2000, 2000–2006, 2007–2010, 2011–2012, 2013, 2014, 2015, 2016, 2017, 2018, 2019, 2020, 2021, 2022. Years prior to 2013 were binned into intervals in such a way that each interval contained at least 7000 complete checklists. This was necessary considering the low volume of information available annually on eBird before 2013 (SoIB 2023b). Month information was reorganized into four seasons to include in analyses: Winter (December–February); Summer (March–May); Monsoon (June–August); and Autumn (September–November).

Finally, we overlaid the spatial grid data onto the eBird data such that every observation would be assigned to a grid cell of each edge resolution (5 km, 25 km, 50 km, 100 km, 200 km). To exclude pelagic checklists (from the ocean), we excluded all checklists in our India dataset that were located outside a 1 km buffer around the country’s boundaries. Thus, we excluded pelagic checklists but retained intertidal checklists.

##### Removing vagrant records

A key step in the SoIB pipeline is to identify and exclude records of vagrants so that local abundance is estimated only within the expected range of a species. With the amount of birdwatching today, vagrants are being found with increasing regularity all over the country. A species prone to vagrancy may have therefore been detected often at multiple sites that do not form a part of its regular range. To prevent such vagrant records from confounding expected patterns of the species of interest, they must be excluded from estimates of abundance and range size (Magurran and Henderson 2003). We assumed that only migratory species would have vagrant records that need to be excluded, and therefore did not consider records of species categorized as ‘Resident’, ‘Resident & Altitudinal Migrant’, ‘Resident & Local Migrant’, “Resident & Localized Summer Migrant”, “Altitudinal Migrant”, or “Resident (Extirpated)” for exclusion.

We used the following procedure. For each month of the year in each 200 km cell, we isolated records of migratory species that had been reported in three or fewer years. Of these cell-month-species combinations, we isolated those that have occurred at least once from 2015 onwards (period for current trends), and removed them from our dataset. Given that each cell may only have been sampled a few times before 2015, three or fewer historical records in a cell does not indicate vagrancy - the species may instead have been regular but not detected in multiple years. On the other hand, the country has been spatially well sampled in recent years and the rare presence of a species in a cell is likely to indicate true vagrancy. After excluding vagrants, complete checklists continue to contain complete information about the presence/absence of every ‘regular’ species, therefore not violating the premise of completeness.

##### Curating eBird data

Data on eBird are subject to rigorous quality checks before they become available for public view and download. A team of over 200 editors curate data quality in India, in addition to the entire community of eBirders who in this process through mechanisms built into eBird, and regional discussion forums outside eBird. Public data nevertheless still contain inaccuracies. Such inaccuracies can be reduced by applying a set of further quality checks on the data.

##### Removing observations unsuitable for analyses

We identified and removed quality-flagged data by:

1. combining data of 17 sets of similar species whose separation are particularly challenging for birdwatchers (**Table S1**).
2. removing observations that were marked by the eBird data review process as not suitable for public output, but were still appearing in the output.
3. removing observations marked as ‘escapee’, a tag used to identify observations of birds that may have escaped from captivity.
4. removing a set of identifiable mistakes in the data (observations out of documented range, checklists plotted in the wrong location, etc.) that crept in even after curation by eBird editors.
5. removing any travelling checklist where the distance specified was >50 km.

##### Reviewing checklist completeness

Birdwatchers who use eBird have varying degrees of familiarity with the platform. Consequently, the dataset contains errors, including those that involve incorrect assignment of protocol and other metadata. We must account for observers erroneously marking a checklist as complete because only truly complete checklists (those that report all species detected) can be used to calculate reporting frequencies. To minimize usage of checklists erroneously marked complete, we relabeled as incomplete those checklists that met any of the following criteria:

1. Protocol was incidental, indicating that birdwatching may not have been the primary focus
2. Distance travelled was greater than 10 km
3. Three or fewer species were reported but the list lacked information on duration
4. Travelling speed (calculated from list metadata) was greater than 20 kmph, indicating compromised detection
5. Duration was less than three minutes
6. Three or fewer species were reported at a rate of less than two species per hour, again suggesting the list was not complete
7. Started at or after 2000 hrs and ended at or before 0400 hrs, because the likelihood of detecting a species drastically changes in the night. This ensured that nocturnal lists did not qualify for LTC and CAT analysis, but still provided distribution information for the species reported.

##### Selecting species and checklists

This section describes how species were deemed to qualify for each analysis based on the amount of available data, and how checklists were selected to use in the analyses of each species.

##### Selecting species

We restricted our focus to the taxonomic level of species, and reports of subspecies or subspecies groups were assigned to the parent species. Of all species reported from India on eBird, we selected a total of 942 species by applying a number of rules. We limited our pool to diurnal species (active during the day) because birdwatching is largely a diurnal activity and nocturnal species are very infrequently detected during the day. From this pool, we selected a species if it qualified through any of three rules:

1. The species had been reported in more than four 200 km grid cells, and had been detected in at least 15 locations in each of the 14 year-intervals. These species qualified for LTC analysis (523 qualifying species). We followed the same selection protocol for CAT analysis but with the year-interval criterion reducing to the eight years from 2015 to 2022 (643 qualifying species).
2. The species is endemic to the Indian Subcontinent or to biogeographical regions therein (109 species).
3. India constitutes a large and/or important part of the species’ breeding or wintering ranges (190 species; see 1. Species attributes, **Fig. 1**).

This procedure to select species for LTC and CAT analyses required species to occur in four 200 km cells, and is therefore only applicable to wide-ranging species. Several species, however, have small ranges but substantial data. Such restricted species sometimes have sufficient long-term data for trend estimation through a different analytical methodology. Of the total species selected (942), we identified a set of range-restricted (detected in four or fewer 200 km grid cells) resident species that had not qualified for LTC and CAT analyses. Only resident species— those with a migratory status of “Resident”, “Resident & Altitudinal Migrant”, “Resident & Local Migrant”, “Resident & Localized Summer Migrant”, “Altitudinal Migrant”, or “Resident (Extirpated)” —were considered. From these, we selected species whose range (in units of 25 km grid cells) contained at least 50 complete checklists in each year-interval. Note that the restricted species list by definition could only contain species that were already part of the 942 but had not qualified for LTC and CAT analyses. A total of 669 species finally qualified for LTC and CAT analyses out of the 942. All 942 species were selected for DRS analysis.

##### Constraining data for a species

For each of the 942 bird species, its LTC, CAT, and DRS had to be estimated using data only from within its broad spatial and temporal occurrence range. This is an important step because the three metrics would otherwise be diluted by the inclusion of information from a location or season where the species cannot be present. Many analyses of structured data (Sauer et al. 2013, North American Bird Conservation Initiative 2014, BirdLife Australia 2015a, b) follow this method, which ensures that local abundances are used to estimate abundance metrics. To restrict data to within the occurrence range of a species, we included, for each month, data from only those 100 km grid cells where the species had been reported. We selected 100 km over 200 km because many regions unsuitable for a species may fall within a 200 km grid cell.

##### Reducing spatial non-independence of checklists

A significant source of bias in any semi-structured data is the overrepresentation of some birdwatching sites (‘localities’ in eBird) in the data because they are heavily frequented by birdwatchers. This leads to pseudoreplication in analyses, and runs the risk of certain sites disproportionately influencing results and inferences. In addition, if the relative representation of birdwatching sites in the data changes over years, inferences for any year are no longer comparable with inferences for every other year. To mitigate this, we spatially subsampled the data so that only a single complete checklist was randomly picked from each site for each month and year-interval, thereby producing a smaller dataset that has equal representation from each site for every month and year-interval. We then repeated this process 1,000 times to produce 1,000 simulated datasets that together represented greater variation from within each site. These filtered datasets were generated for use in the analyses of LTC and CAT, but not of DRS.

### Estimating long-term change (LTC) and current annual trend (CAT)

We selected the entire period before the year 2000 as the baseline against which the changes in abundance were compared in the long term. The year 2000 was chosen as the threshold year because it was the earliest period for which aggregated data were found to be adequate for abundance (frequency) estimation. The LTC metric was defined as the percentage change in the frequency of reporting between the year 2022 and the year-interval pre-2000. We selected 2015 as the baseline for the CAT because we found that annual patterns of eBird usage in India are most comparable in the years starting from 2015, probably because eBird was first used in India in 2013 and patterns of eBird usage took some time to develop before settling. The CAT is defined as the mean percentage annual change in frequency during the eight years from 2015 to 2022. Given the large amount of data available for this period and comparable eBirding patterns, we are especially confident about the CAT metric. Consequently, this confidence also reflects in the greater relative weight given to the CAT over the LTC when making species-level prioritization decisions (See **Fig. 1**, 4.).

#### Modelling frequency of reporting

This section describes the steps followed to build and execute statistical models to estimate frequency of reporting for each species during each year-interval.

#### Running single species models

We considered each checklist from the curated dataset to be an independent sampling unit and modelled detection/non-detection (the response variable) for each species using binomial Generalized Linear Mixed-effects Models (Bird et al. 2014, Walker and Taylor 2017, Horns et al. 2018). Although data on non-detection are not present in eBird data, such data are implicit for all species absent in complete checklists. The primary explanatory variable (fixed effect) was the year-interval (as a factor) because our main objective was to infer trends over years. We did not include this as a continuous variable because we did not want to force a monotonically decreasing or increasing trend over years, and instead wanted to be able to detect fluctuations at a yearly interval. Other explanatory variables (fixed effects) were season (factor), and the interaction between season and list length (whole number). We included these two variables to statistically control for the effects of seasonality and birdwatching effort on detection.

Random effects were specified as a nested 25 km–within–100 km grid cell structure to smoothen spatial bias and account for spatial autocorrelation (**Eq. 1**). We specified fewer random effects for species restricted in range (see **Fig. 1**, 1., Selecting species and checklists, Selecting species) in the following way: 1) a 25 km grid cell random effect was included for any restricted species that was reported in eight or more 25 km grid cells (**Eq. 2**); 2) no random effect was specified for any restricted species that was reported in seven or fewer grid cells (**Eq. 3**). We also considered including observer identity as a random effect to account for varying observer ability (Sauer et al. 1994, Link and Sauer 1998, Kelling et al. 2015, Johnston et al. 2018), but decided against it for two reasons: 1) observer random effects appeared to make little to no difference to parameter estimates from exploratory analyses but made the models data-hungry; 2) observer abilities change with time, and we would have had to effectively incorporate a separate effect within each year-interval (a random slope), which would be computationally intensive.

We used a complementary log-log link function to transform our binomial response as it approaches asymptotes more gradually than a logit link function and is therefore most appropriate for the majority of species that are either very rare or very common. We used the functions glm() and glmer() from the *lme4* package (Bates et al. 2015) in R 4.2.3 (R Core Team 2023) to fit these models and obtain predictions. We modelled the probability of reporting a species (P) in the following manner (‘j’ represents every checklist):

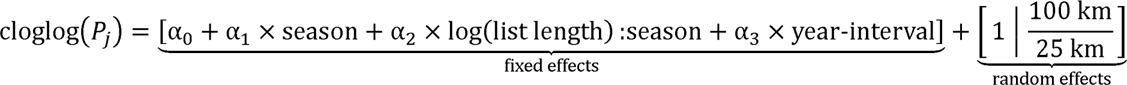

**Eq. 1**: Species that are widespread.

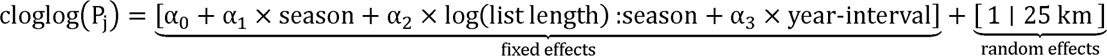

**Eq. 2**: Species that are moderately restricted (reported in eight or more 25 km grid cells).

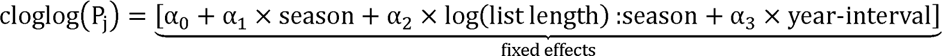

**Eq. 3**: Species that are highly restricted (reported in seven or fewer 25 km grid cells).

For the mixed-effects models, we used the function predictInterval() from the package *merTools* (Knowles et al. 2023) to obtain the mean reporting frequency and associated standard error (SE) of a species for a list of median length in each season for each year-interval. For the fixed-effects models, we obtained predictions from the base function glm::predict(). In both cases, means and SEs were averaged across the four seasons to get a single mean and SE (individual errors were propagated) for the species in each year-interval. This process was repeated for all 1,000 randomly generated datasets for all qualifying species, effectively bootstrapping and providing a single file with 1,000 different mean and SE values for each year-interval for each species. Note that these values of reporting frequency were still on the transformed (Gaussian) scale and were later back-transformed to calculate LTC and CAT (see headings titled ‘Calculation’), and the associated confidence intervals.

#### Model confidence checks

We needed to ensure that any species selected for LTC and CAT analyses were widely distributed within 25 km cells, thereby reducing pseudoreplication and spatial autocorrelation. We therefore excluded from LTC and CAT calculations any species: 1) that had a mean of less than eight 5 km subcells sampled with at least one complete checklist, out of a maximum of 25 cells, across all 25 km grid cells where the species occurred; or 2) for which the variation in the proportional coverage of 5 km subcells within each 25 km cell was so high that the CI:mean ratio was greater than 0.25. This step was not applicable to species restricted to the islands because many 5 km subcells within a 25 km grid cell may be entirely located in the ocean and therefore be inherently out of reach. Next, we needed to ensure greater comparability between the frequencies estimated for these years. We therefore excluded any species whose average sampled range (25 km grid cells with complete lists) between 2015 and 2022 was less than 60% of its sampled range in 2022.

Additionally, we ran some checks to reduce ‘problematic’ predictions of trends in any of the 1,000 simulations, either due to a species not having sufficient data or because of model convergence issues. For species with LTC estimates, we removed predictions for every simulation where the transformed SE for any year-interval before 2015 was NA (not estimated) or where the transformed SE was greater than the absolute value of the mean (an arbitrary high threshold, for greater caution). After this process, if less than 500 predictions were remaining (out of 1000) for a species, its LTC metric was not calculated. For species with only CAT estimates (2015 onwards), we followed the same process but added the additional criterion of the absolute value of the predicted estimate for a year-interval not exceeding 100, indicating near 0 or 1 (back-transformed) frequency values and a problem with convergence.

#### Predicting values

We averaged predicted values across simulations (up to 1,000) for each species and each year-interval. This averaging was done on the transformed (Gaussian) values, where SEs were assumed to be symmetrical about the mean. We obtained a single mean value for each species and each year-interval, and calculated SE by allowing errors to propagate. This estimate referred to later as “transformed frequency of reporting” is used in subsequent calculations.

#### Estimating LTC and CAT

##### Calculation of LTC

We back-transformed the transformed reporting frequency estimates using the clogloglink() function to get the actual frequency of reporting (ranging from 0 to 1) as well as lower and upper 95% confidence intervals (using a ×1.96 conversion on the transformed SE estimates). We proceeded to standardize the actual frequency of reporting for every species so that an inference of change could be made relative to the first year-interval, which would then be comparable across species, unlike the raw reporting frequencies. We achieved this standardization by dividing the estimate for every year-interval by the pre-2000 estimate, then multiplying by 100 and subtracting 100. Standardized frequency of reporting is referred to as “abundance-index”, henceforth. LTC is defined as the abundance-index for the year 2022.

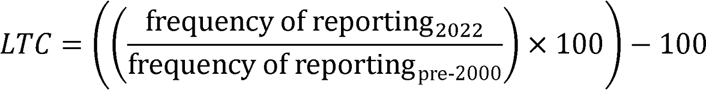

**Eq. 4**: Calculation of long-term change (LTC).

To calculate the uncertainty associated with each abundance-index for each species, we combined the uncertainty in every reporting frequency estimate with the uncertainty in the reporting frequency estimate for the pre-2000 period. A very imprecise pre-2000 estimate would therefore result in a lack of confidence in the abundance-index for every subsequent year-interval. We achieved the calculation of uncertainty by repeating the abundance-index calculation for each year interval 1000 times using reporting frequency values sampled from the Gaussian distribution described from the mean and SE of the respective transformed reporting frequencies. We estimated 95% CIs from the 1000 abundance-index estimates for each year-interval for each species.

##### Categorization of LTC

We drew from published State of Birds reports that used five-way assessment schemes to categorize long-term metrics (Blancher et al. 2009, North American Bird Conservation Initiative 2012, Hayhow et al. 2017). In addition to these, we included two categorizations that are not often part of State of Birds assessments: ‘Insufficient Data’ and ‘Trend Inconclusive’. The use of ‘Trend Inconclusive’ was to safeguard against making inferences using any estimates that were associated with high uncertainty. A species was ‘Stable’ if it did not qualify for decline or increase, nor was the trend inconclusive. Stable here therefore implies that the trend is ‘no different from stable’, a deliberately conservative rule of thumb. Criteria used to place each species in LTC categories are explained in **Fig. 3**. The decisions are hierarchical, and species were classified in the order of the specified rules (top to bottom). Specified thresholds apply to lower confidence limits for increases and upper confidence limits for declines.

**Fig. 3:**
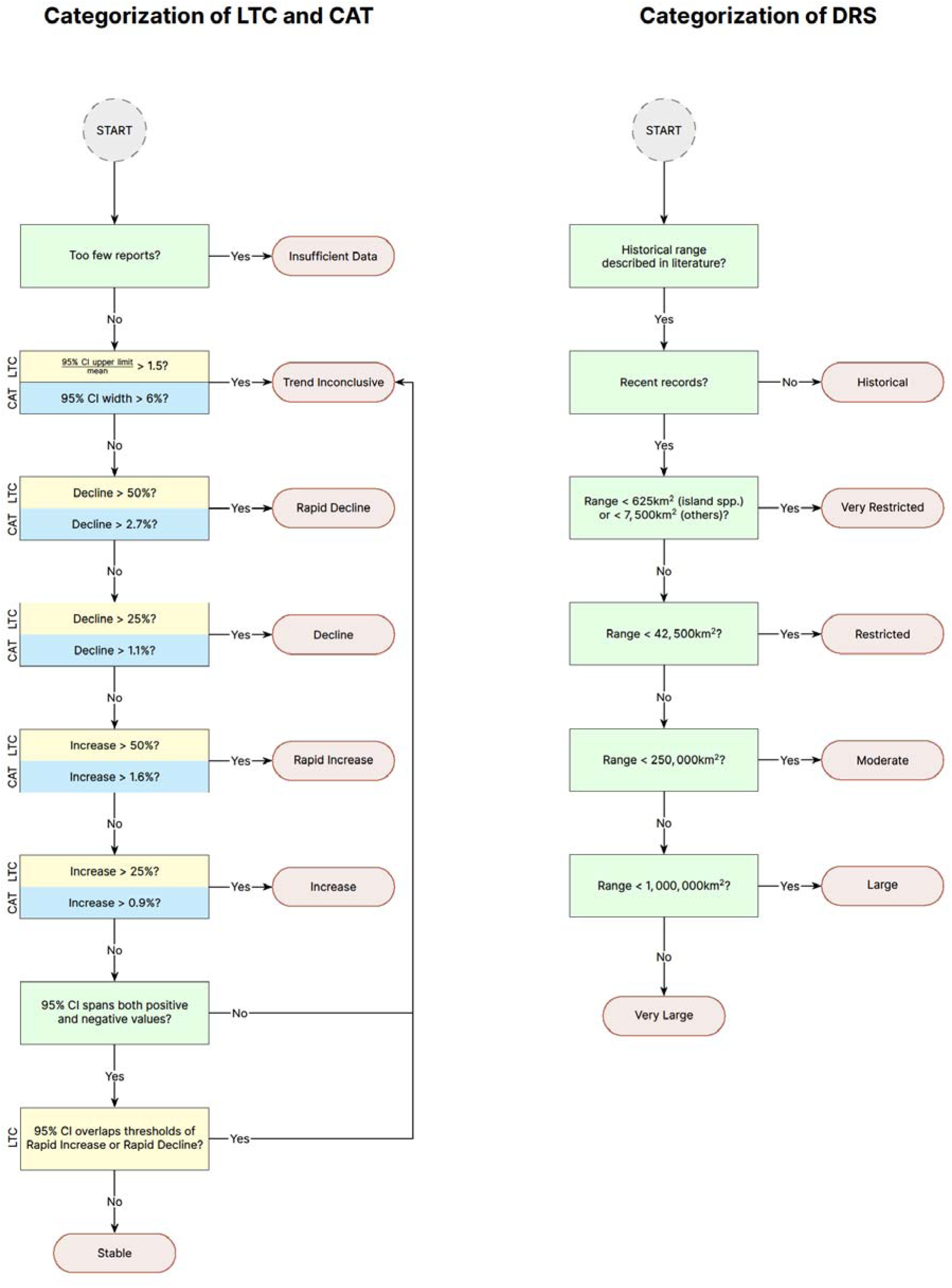
Decision tree to categorize LTC, CAT, and DRS.

##### Sensitivity check of LTC

We took the results through a sensitivity filter to ensure that the LTC is robust and less likely to change significantly due to statistical randomness in the algorithms when the calculation (which includes simulations) is repeated. We repeated the process of calculating the LTC five times, thereby repeating the process used to calculate uncertainty, which involved random sampling from Gaussian distributions with means and SDs corresponding to model predictions. For each species, we recategorized the LTC based on each of the five LTC calculations. We compared categories from these five repetitions with the original categories (a total of six) and changed the original category hierarchically to: 1) Trend Inconclusive if a species was placed in more than two categories; 2) Increase if the simulations produced both Rapid Increase and Increase classifications; 3) Decline if the simulations produced both Rapid Decline and Decline classifications; 4) Trend Inconclusive if the species was placed in two contradictory categories (e.g., Stable and Decline).

##### Calculation of CAT

For each species in each year-interval from 2015 to 2022, we simulated 1,000 back-transformed values by sampling from the distribution of transformed reporting frequency predictions derived from the models. We assumed that the relationship between reporting frequency and time is near-linear for the short duration of eight years (Buckland et al. 2005). We randomly sampled a frequency value for each species and year-interval from the 1,000 simulated values, and ran a linear model with these eight sampled values against year-intervals (a continuous variable this time, 2015–2022). We calculated a mean reporting frequency for each year-interval (and the associated propagated SE) by resampling the eight values 1000 times and running linear regressions, and finally averaging across the 1000 predictions obtained for each year-interval. To estimate CAT, defined as the percentage annual change in reporting frequency over the eight year period, we selected the year-intervals 2015 and 2016 (the highest uncertainty years so that the results would be most conservative), and calculated the slope of the regression proportional to the 2015 reporting frequency estimate (standardized to a percentage). We did not directly fit simple linear models onto the raw data with a continuous predictor that spanned the last eight years, because our response (detection/non-detection) is binary and requires a non-linear transformation even if the transformed relationship is linear.

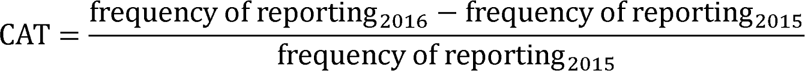

**Eq. 5**: Calculation of current annual trend (CAT) from the predictions of a linear regression that uses predictions from the main model as input.

##### Categorization of CAT

CAT criteria are defined such that the thresholds for annual change (**Fig. 3**) compounded over 30 years (from the pre-2000 median year 1992 to 2022), approximately equals our long-term thresholds, therefore making long-term and current categories comparable. The categories and the process of categorization are otherwise as described in the LTC categorization section.

##### Sensitivity check of CAT

We took the results through a sensitivity filter to ensure that the CAT is robust and not likely to change substantially if estimates from any one of the eight past years were dropped from the calculation. This is to reduce instances where a single year disproportionately drives estimates of the CAT. We recalculated the CAT eight more times for each species, each time dropping one of the eight years that were considered in the calculation. We compared the nine such classifications (one original and eight sensitivity reclassifications for each species), and changed the original category hierarchically to: 1) Trend Inconclusive if a species had at least one pair of qualitatively opposite (increase vs decrease) categorizations; 2) Trend Inconclusive if a species had four or more Trend Inconclusive classifications out of the nine; 3) Increase if it was originally Rapid Increase but at least one other reclassification produced an Increase classification; and 4) Decline if it was originally Rapid Decline but at least one other reclassification produced a Decline classification.

### Estimating distribution range size (DRS)

The use of eBird data provides the unique possibility of quantifying species range sizes. We decided to calculate DRS based on the actual occupied range and not the modelled range, because species distribution models provide predictions even for regions (grid cells) where the species may not occur. We used an occupancy framework to account for imperfect detection in grid cells where the likelihood of occurrence was high, i.e., in the grid cells neighbouring those where the species had been detected.

#### Modelling occupancy

Our objective was to compute the probability of occupancy of every 25 km grid cell for every selected species, but only from 2018 to 2022, so that the estimated DRS is current. This extant range would include the area where the species had been detected, as well as the probable neighbouring area it is expected to occupy where it had not yet been detected. Any species with more than one seasonal distribution range within the country would require more than one occupancy estimated. We therefore classified species into one or more of the following categories based on the hierarchical rules in **Table S3**: ‘Resident’ (R), ‘Migrant Summer’ (MS), ‘Migrant Winter’ (MW), ‘Migrant Passage’ (MP), and ‘Migrant’ (M).

#### Running single-species models

We assumed that the probability of occupancy of a species to be 1 in those grid cells where it was detected. To estimate the probability of occupancy of a species in neighbouring cells where it was not detected, we used the R package *unmarked* (Fiske and Chandler 2011) and modelled species occupancy in their extant range (past five years, i.e., 2018–2022) while accounting for imperfect detection (Kéry et al. 2010). For each cell, we calculated one metric to be used as a site covariate—the proportion of occupied neighbouring 25 km grid cells out of eight—so that unsampled cells can have some estimated finite probability of occupancy in the neighbourhood of occupied cells (Altwegg and Nichols 2019). Season and list length were the checklist-level covariates considered. To reduce violations of closure assumptions, we ran separate models spanning different seasons for migratory and resident species (Altwegg and Nichols 2019). We modelled occupancy (ψ) and detection probability (P) in the following manner (‘i’ represents every 25 km edge cell and ‘j’ represents every checklist within a cell):

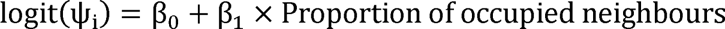

**Eq. 6**: Occupancy.

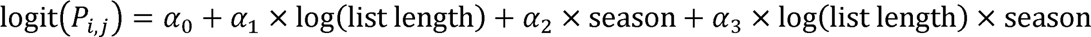

**Eq. 7**: Detection probability of species that are resident.

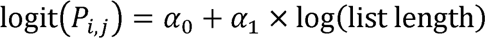

**Eq. 8**: Detection probability of species that are migratory.

We ran the appropriate model for each species for each season (where applicable) to predict the occupancy probability in each unoccupied neighbouring cell based on the proportion of its occupied neighbours. We also predicted detection probability (P) for each checklist in a cell, and then subtracted that from 1 to get the probability of the species not being detected in each checklist. By multiplying non-detection probabilities across all checklists within a cell, we calculated the probability of non-detection of the species within the cell. SEs were propagated while calculating the product (see Michigan State University 2019). Finally, we multiplied the probability of non-detection of a species in a cell with the predicted occupancy in that cell to calculate a more informed probability of the species occupying that cell. SEs were again propagated as mentioned previously. Note that the probability of a species occupying a cell where it had been reported is set as 1, and was set as 0 for every cell aside from these and their immediate neighbours. These cell-specific probability values were later used together with the area of a grid cell to estimate DRS for each species.

#### Predicting values

We calculated the expected range size associated with a 25 km grid cell of a species separately for each seasonal distribution (R, M, MS, MW, MP). This was calculated as the product of the area of a cell (in km^2^) with its predicted occupancy (1 for cells with confirmed presence, 0 for ‘non-detection’ cells with no confirmed neighbour, and non-zero (0–1 exclusive) for ‘non-detection’ cells with at least one confirmed neighbour).

#### Estimating DRS

##### Calculation of DRS

Regionwide range size was then estimated by summing across expected cell-level range sizes while propagating SEs derived from the occupancy model. This yielded a number (often large) that can be interpreted as the total area within the country in which the species is currently likely to occur. When a species had more than one seasonal distribution, we picked the largest of the estimated seasonal range sizes as the DRS for that species.

##### Categorization of DRS

We adopted a five-way assessment scheme to categorize DRS (**Fig. 3**). Species restricted to islands within India have different rules for categorization as Very Restricted because they inherently have small range sizes. Any species selected for SoIB that did not have any reports during the past five years were categorized as ‘Historical’.

### Assigning categories of conservation priority

A species was assigned a category of Conservation Priority based on the classifications of the three metrics LTC, CAT, and DRS. For example, a species that had its LTC and CAT classified as Rapid Decline was categorized as ‘High’ Conservation Priority. The set of rules for this categorisation are comprehensively outlined in the Methods section of the SoIB (2023b). In cases where the trend classifications were either Insufficient Data or Trend Inconclusive, the species’ IUCN Red List status also figured in the priority categorisation.

### Code

All the code associated with the SoIB 2023 analyses and outputs is housed in a GitHub repository (stateofindiasbirds 2024).

## Results

### Summary results

All metrics taken together, 178 species were identified as of High Conservation Priority, 323 as Moderate, and 441 as Low. Of the 178 High Priority species, 94 were categorised based only on information about either or both of LTC and CAT, together with DRS, 45 based on DRS alone, and 39 after factoring in their IUCN Red List status. The High Priority species include 90 that are Least Concern on the global IUCN Red List, and a further 17 that are Near Threatened. Among species that are globally threatened on the IUCN Red List, two Vulnerable (Greater Spotted Eagle and Great Hornbill) and one Endangered species (Steppe Eagle) were identified as of Low Priority (**Table 1**).

**Table 1:** Cross-tabulation of IUCN Red List status with SoIB priority status.

|  | High | Moderate | Low | Total |
| --- | --- | --- | --- | --- |
| Critically Endangered | 14 | 0 | 0 | 14 |
| Endangered | 15 | 0 | 1 | 16 |
| Vulnerable | 42 | 8 | 2 | 52 |
| Near Threatened | 17 | 39 | 11 | 67 |
| Least Concern | 90 | 269 | 422 | 781 |
| Not Recognised | 0 | 7 | 5 | 12 |
| Total | 178 | 323 | 441 | 942 |

Of the 523 species with LTC estimates, 338 had robust trends (not Trend Inconclusive). Of these, 98 declined in abundance by more than 50% in the long term, meeting the criterion for Rapid Decline. Of the 643 species with CAT estimates, 359 had robust trends. Of these, 142 showed evidence of decline, and 189 were stable. Of the 942 species with DRS estimates, 266 were Restricted or Very Restricted (**Table 2**).

**Table 2:** Break-up of species categorizations in the three SoIB metrics.

| **Category: LTC and CAT** | **Species: LTC** | **Species: CAT** | **Category: DRS** | **Species: DRS** |
| --- | --- | --- | --- | --- |
| Rapid Decline | 98 | 64 | Historical | 4 |
| Decline | 106 | 78 | Very Restricted | 46 |
| Stable | 98 | 189 | Restricted | 220 |
| Increase | 19 | 17 | Moderate | 363 |
| Rapid Increase | 17 | 11 | Large | 178 |
| Trend Inconclusive | 185 | 284 | Very Large | 131 |
| Insufficient Data | 419 | 299 |  |  |

Species with restricted range sizes typically did not have sufficient data to estimate trends with high confidence (**Table 3**). Only 2% of Very Restricted and 1% of Restricted species had statistically robust LTC estimates, all of which were declines. None of the Very Restricted species had CAT estimates within accepted limits of confidence intervals, but 5% of Restricted species did, all of which were declines. Around 40% of species with Moderate range size had LTC and CAT estimates within accepted limits of confidence intervals. A greater proportion of Large range species were categorized as in Decline or Rapid Decline (46% for LTC and 29% for CAT) than Very Large range species (23% for LTC and 19% for CAT). Conversely, more Very Large range species showed an Increase or Rapid Increase, than did Large range species.

**Table 3:** Cross-tabulation of distribution range size categories with each trend category.

| **DRS** | **LTC** |  |  |  |  |  |  |
| --- | --- | --- | --- | --- | --- | --- | --- |
|  | Insufficient<br>Data | Trend<br>Inconclusive | Rapid<br>Decline | Decline | Stable | Increase | Rapid<br>Increase |
| Very<br>Restricted | 96 | 2 | 2 | 0 | 0 | 0 | 0 |
| Restricted | 94 | 5 | 0 | 1 | 0 | 0 | 0 |
| Moderate | 42 | 20 | 12 | 12 | 12 | 1 | 1 |
| Large | 5 | 30 | 23 | 23 | 16 | 2 | 1 |
| Very Large | 2 | 37 | 15 | 8 | 21 | 8 | 8 |
|  | **CAT** |  |  |  |  |  |  |
| Very<br>Restricted | 91 | 9 | 0 | 0 | 0 | 0 | 0 |
| Restricted | 81 | 14 | 4 | 1 | 0 | 0 | 0 |
| Moderate | 18 | 44 | 6 | 9 | 21 | 1 | 0 |
| Large | 3 | 30 | 13 | 16 | 33 | 3 | 2 |
| Very Large | 2 | 27 | 8 | 11 | 40 | 6 | 5 |

### Species with the steepest increases

Glossy Ibis and Indian Peafowl are among the few species that have increased tremendously in the long term, by 136% and 124% (lower CIs) respectively (**Table 4**). Glossy Ibis, however, has a Stable CAT whereas Indian Peafowl continued to show a current Rapid Increase of 6% per year.

**Table 4:**
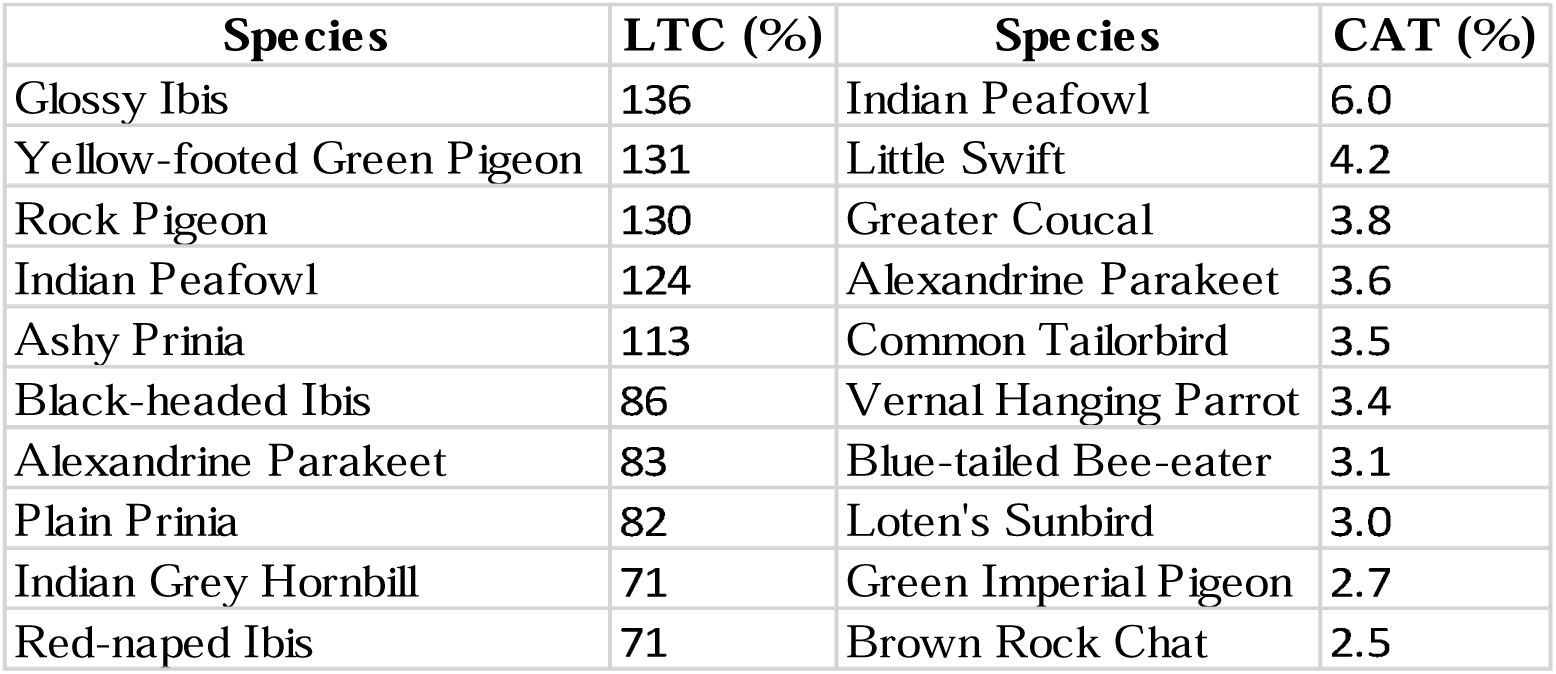
The ten species with the greatest long-term change (LTC) and current annual trend (CAT) arranged in descending order of magnitude of their lower CIs. Species with LTC or CAT categorized as Trend Inconclusive are excluded here.

| Species | LTC (%) | Species | CAT (%) |
| --- | --- | --- | --- |
| Glossy Ibis | 136 | Indian Peafowl | 6.0 |
| Yellow-footed Green Pigeon | 131 | Little Swift | 4.2 |
| Rock Pigeon | 130 | Greater Coucal | 3.8 |
| Indian Peafowl | 124 | Alexandrine Parakeet | 3.6 |
| Ashy Prinia | 113 | Common Tailorbird | 3.5 |
| Black-headed Ibis | 86 | Vernal Hanging Parrot | 3.4 |
| Alexandrine Parakeet | 83 | Blue-tailed Bee-eater | 3.1 |
| Plain Prinia | 82 | Loten's Sunbird | 3.0 |
| Indian Grey Hornbill | 71 | Green Imperial Pigeon | 2.7 |
| Red-naped Ibis | 71 | Brown Rock Chat | 2.5 |

### Patterns in abundance trends based on diet

Birds that specialize on diets of vertebrate prey, carrion or invertebrates as a composite have faced the steepest long-term declines of more than 25%. Birds that depend on fruits or nectar, in contrast, have remained stable or even shown signs of increase in the long term (**Fig. 4**).

**Fig. 4:**
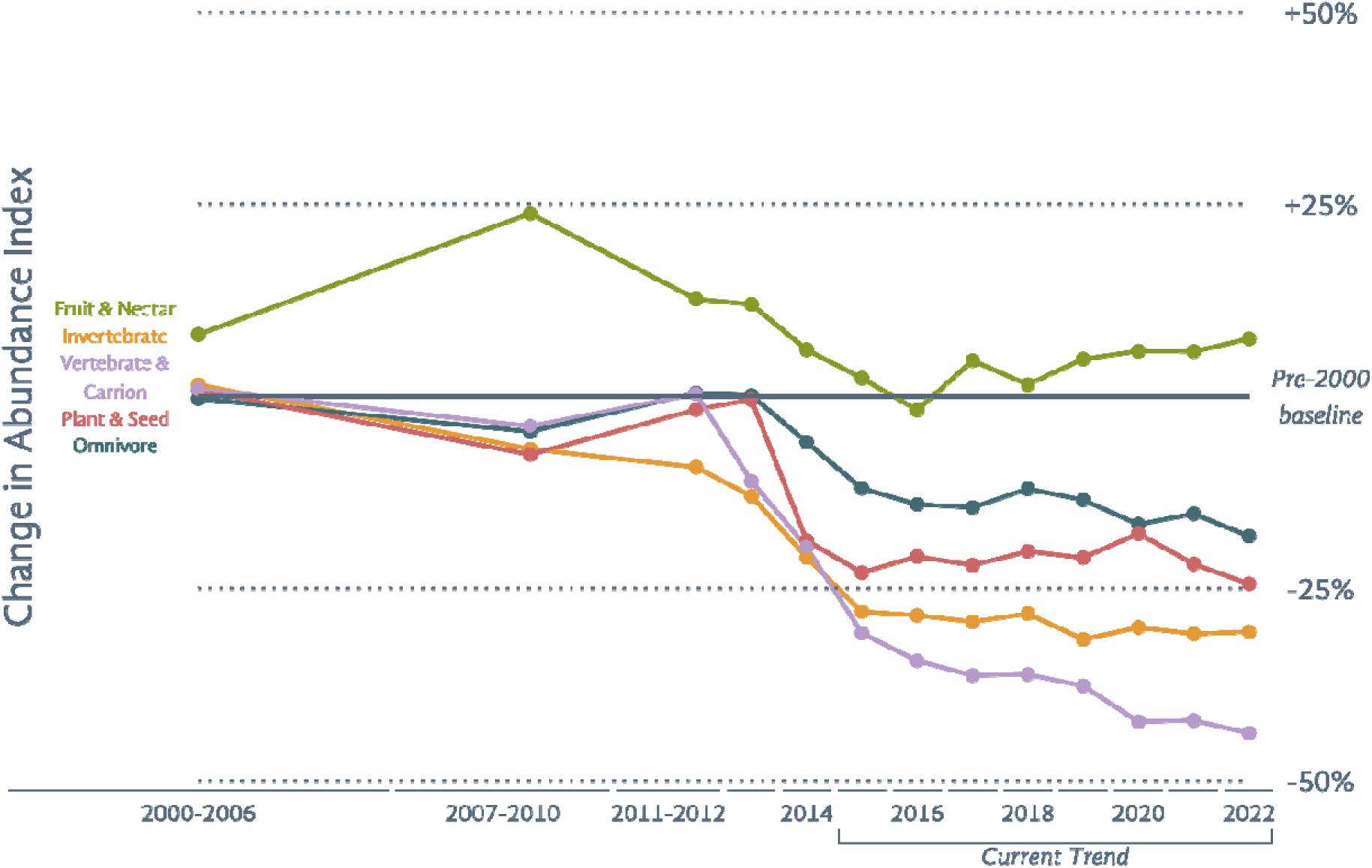
A comparison of the composite trends of birds that belong to five diet groups. Composite trends were calculated by averaging the mean individual trends of several species. Points refer to the proportional change in reporting frequency when compared to the pre-year-2000 baseline. Note that CIs have not been depicted for ease of interpretation. For detailed information about how to interpret this graph and for the lists of constituent species included in the composites, see the Methods section in the SoIB 2023 report (SoIB 2023b).

Among the vertebrate and carrion feeders, most vulture species in India have undergone a Rapid Decline in the long term, except Slender-billed Vulture for which data was insufficient to estimate a trend, and Himalayan Griffon which has faced a Decline. Of the vultures with robust CAT estimates, four species continued to be in Rapid Decline, including Indian Vulture which has been declining most rapidly at 8.7% a year (upper CI limit), and two species in Decline including White-rumped Vulture (**Fig. 5**).

**Fig. 5:**
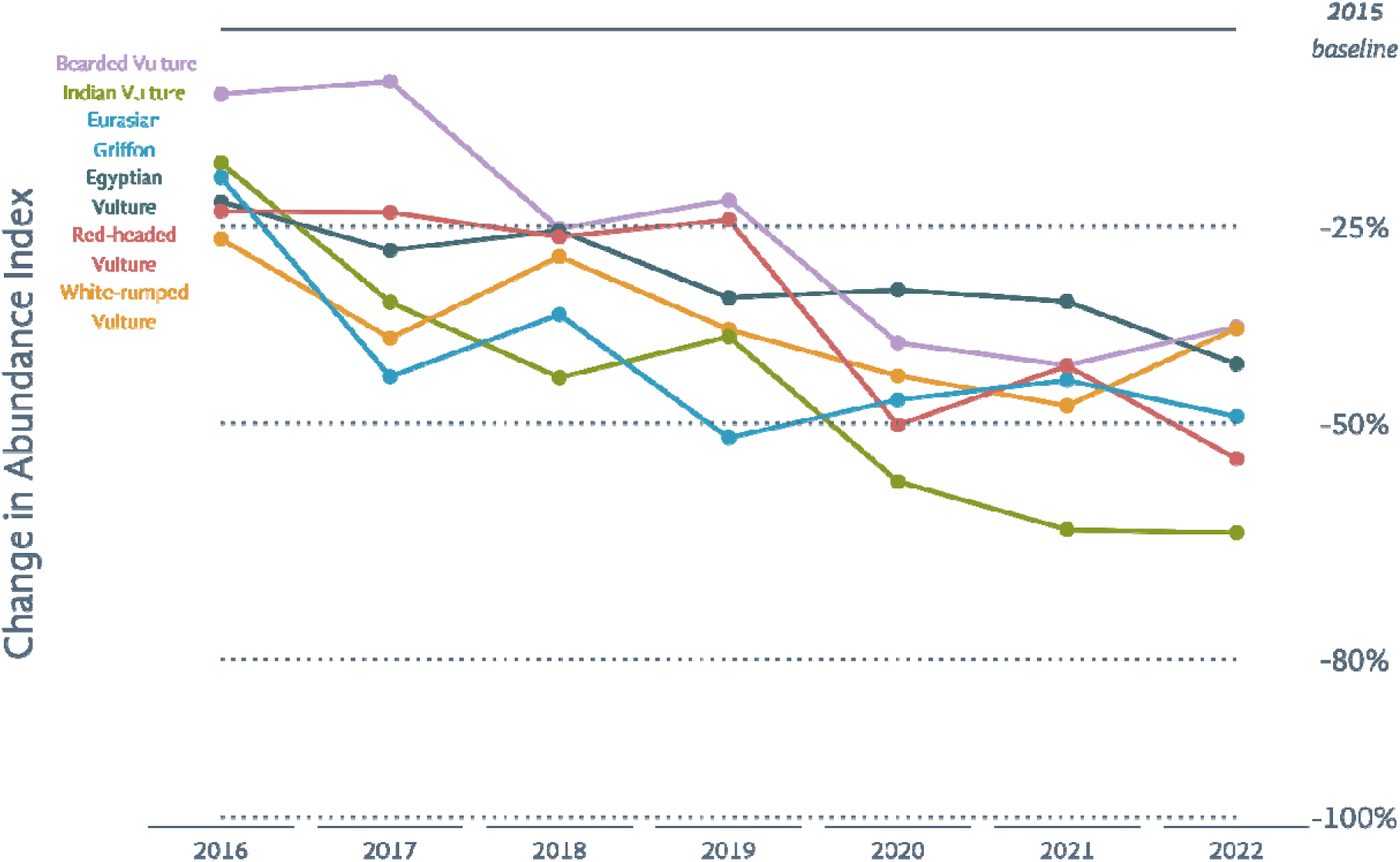
A comparison of current trends (2015–2022) of six vulture species. Points refer to the proportional change in reporting frequency when compared to the 2015 baseline. Note that CIs have not been depicted for ease of interpretation.

Another group of vertebrate feeders that have declined is harriers. All harriers with robust trend estimates—Montagu’s, Pallid, and Western/Eastern Marsh-Harrier—have undergone a Rapid Decline in the long term and have continued to face a ∼3% annual Decline in recent years. Data was sparse for large *Aquila* and *Clanga* eagles, but Tawny Eagle stands out with a Rapid Decline in the long term and a continuing Rapid Decline even today. Steppe, Bonelli’s and Greater Spotted Eagles show Stable current trends. Greater Spotted Eagle has also been Stable in the long term.

### Patterns in abundance trends based on habitat

Birds that specialize on grassland and scrub habitats have declined the most, followed by those that require more generic ‘open habitats’ or wetlands (**Fig. 6**). Among the nine resident and migratory species that specialize on dry grassland and scrub habitats and have robust LTC estimates, seven have undergone a Rapid Decline, and the remaining two a Decline, in the long term. In recent years, however, none of these species are in Rapid Decline although five—Indian Courser (72% long-term decline), Pallid Harrier (72%), Montagu’s Harrier (55%), Great Grey Shrike (83%), Isabelline Wheatear (58%)—are in annual Decline of ∼3%. Laggar Falcon, another dry grassland specialist, did not have sufficient data for LTC analysis but is in Rapid Decline of ∼4% currently.

**Fig. 6:**
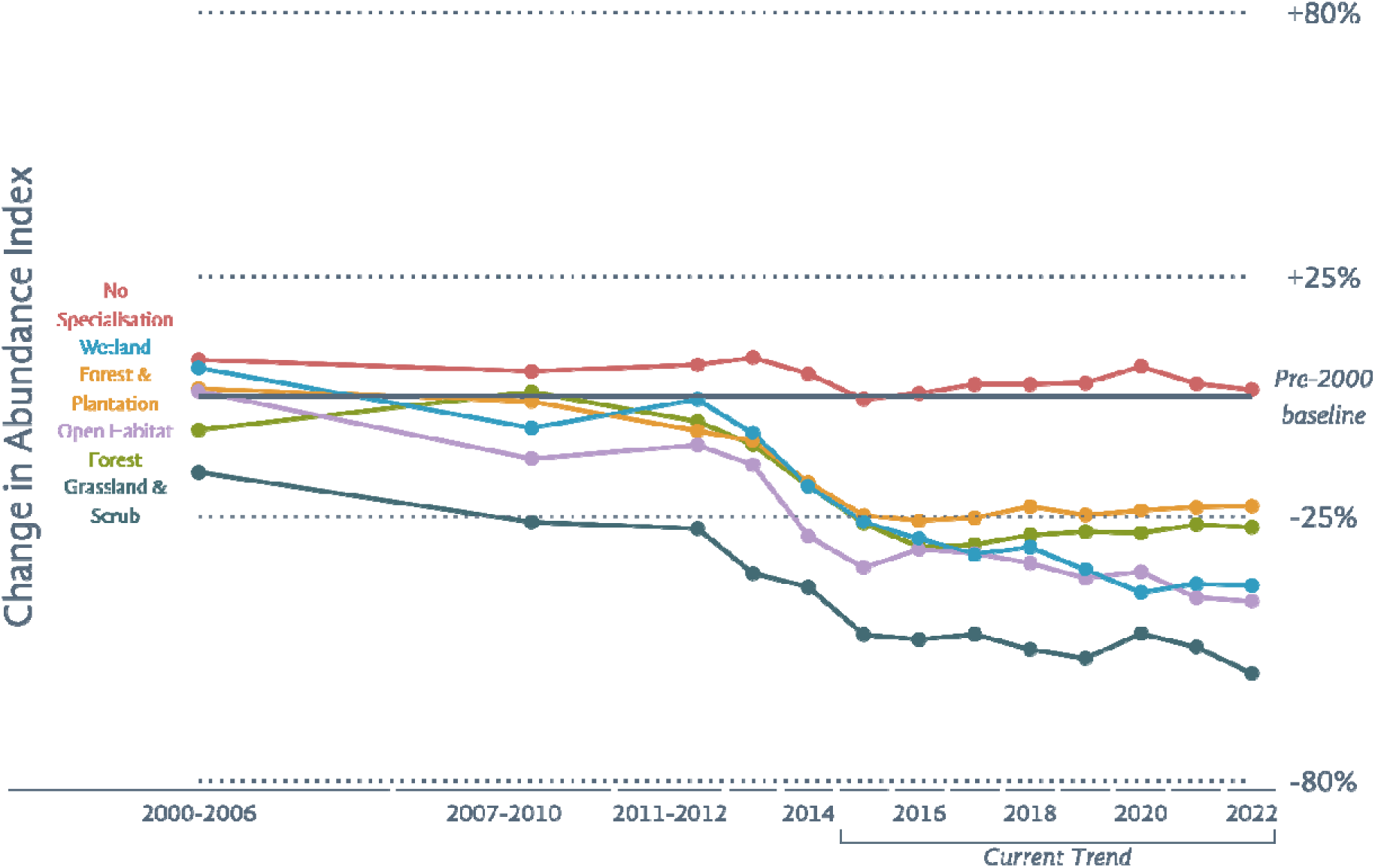
A comparison of the composite trends of birds that belong to six habitat groups. Composite trends were calculated by averaging the mean individual trends of several species. Points refer to the proportional change in reporting frequency when compared to the pre-2000 baseline. Note that CIs have not been depicted for ease of interpretation. For detailed information about how to interpret this graph and for the lists of constituent species included in the composites, see the Methods section in the SoIB 2023 report (SoIB 2023b).

Some groups of wetland birds such as ducks and shorebirds have declined more than others. Of the twelve duck species with robust trends, only Lesser Whistling-Duck has a Stable LTC, while the rest have all declined. Seven of the remaining eleven have undergone a Rapid Decline, with the greatest declines of over 70% experienced by Northern Pintail and Tufted Duck. All ducks that had declined in the past continued to be in Rapid Decline (6 of 11 species) or Decline (4 of 11 species) today, except Red-crested Pochard which shows a Stable current trend.

Other groups of waterbirds show mixed trends. 16 species show a Stable or increasing CAT, while 23 species show a decreasing CAT, of which 10 species are in Rapid Decline, including Greater Flamingo (6% per year), Baillon’s Crake (7%), Sarus Crane (4%), Spot-billed Pelican (4%), and Eurasian Spoonbill (6%). Spot-billed Pelican had shown an increasing trend until c. 2013 before rapidly decreasing to its pre-2000 levels by 2022. Greater Flamingo, Sarus Crane, Eurasian Spoonbill and Black-capped Kingfisher have also shown a Rapid Decline in the long term. Several species are in current Decline such as Lesser Adjutant, Painted Stork, and Pied Kingfisher.

### Patterns in abundance trends based on migratory behaviour

Birds that migrate to India during the winter have declined more than those that are resident (**Fig. 7**).

**Fig. 7:**
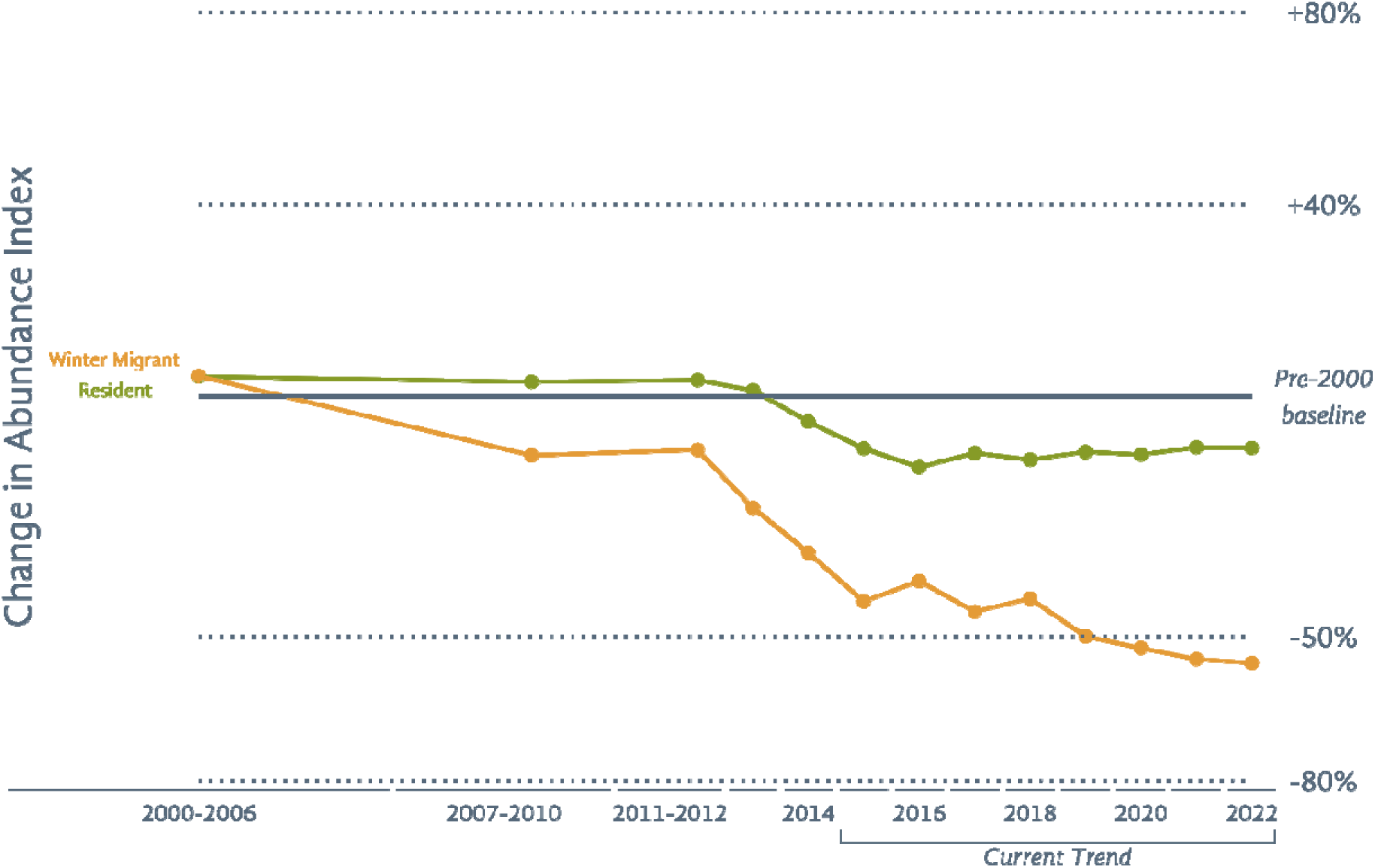
A comparison of the composite trends of Indian resident birds and winter migrant birds. Composite trends were calculated by averaging the mean individual trends of several species. Points refer to the proportional change in reporting frequency when compared to the pre-2000 baseline. Note that CIs have not been depicted for ease of interpretation.

Among all winter migrants to India, Black-capped Kingfisher has declined the most in the long term (86%, lower CI). A large proportion of the species that have declined the most are migratory shorebirds (**Table 5**). The only non-wetland migratory species that feature in the top 10 declining birds (LTC and CAT) are Eurasian Griffon and Pallid Harrier with declines of 79% and 73% in the long term, and Forest Wagtail with a current annual decline of 5.9%.

**Table 5:** The ten species that are winter visitors to India and have the greatest declines in long-term change (LTC) and current annual trend (CAT), arranged in descending order of the absolute magnitude of their lower CIs. Species with LTC or CAT categorized as Trend Inconclusive are excluded here.

| Species | LTC (%) | Species | CAT (%) |
| --- | --- | --- | --- |
| Black-capped Kingfisher | -86 | Sandwich Tern | -8.6 |
| Dunlin | -82 | Eurasian Oystercatcher | -7.1 |
| Eurasian Griffon | -79 | Baillon's Crake | -6.9 |
| Eurasian Curlew | -77 | Dalmatian Pelican | -6.5 |
| Pied Avocet | -77 | Northern Shoveler | -6.0 |
| Curlew Sandpiper | -76 | Forest Wagtail | -5.9 |
| Terek Sandpiper | -75 | Northern Pintail | -5.9 |
| Grey Plover | -73 | Spotted Redshank | -5.6 |
| Tufted Duck | -73 | Ruddy Turnstone | -5.5 |
| Pallid Harrier | -73 | Terek Sandpiper | -5.4 |

### Patterns in abundance trends based on endemism

Species that are endemic to the Western Ghats have declined more than endemics to other regions and non-endemics (**Fig. 8**).

**Fig. 8:**
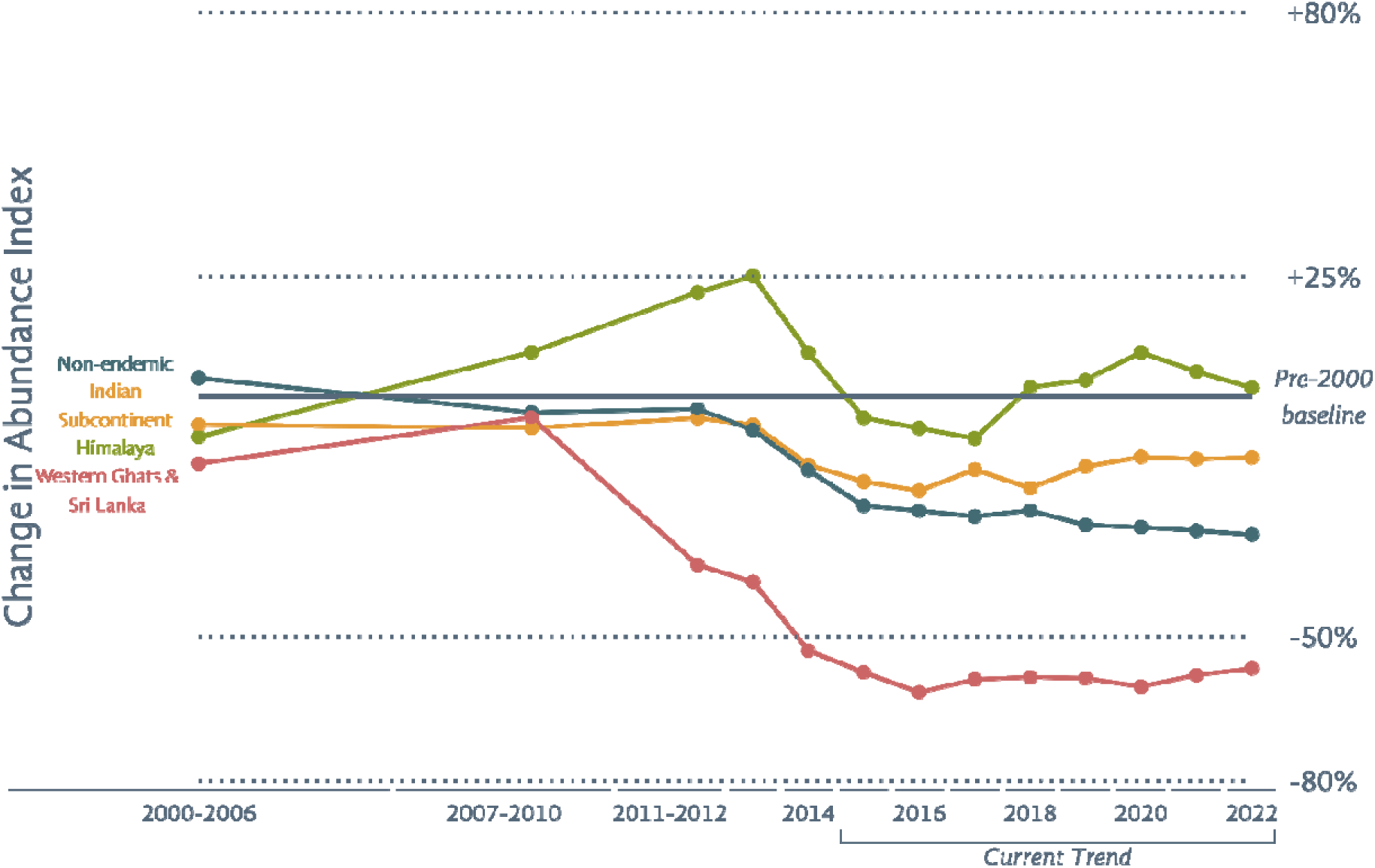
A comparison of the composite trends of birds that are endemic to different regions. Composite trends were calculated by averaging the mean individual trends of several species. Points refer to the proportional change in reporting frequency when compared to the pre-2000 baseline. Note that CIs have not been depicted for ease of interpretation. For detailed information about how to interpret this graph and for the lists of constituent species included in the composites, see the Methods section in the SoIB 2023 report (SoIB 2023b).

## Discussion

### A framework to leverage semi-structured citizen science for conservation

We present a detailed framework to assess the status of birds in a region using semi-structured data uploaded to eBird. Our methodology not only enables vital status assessments in the absence of region-wide systematic monitoring schemes, but also allows larger numbers of species to be assessed when systematic schemes sometimes cover a smaller set of species (e.g., Lin and Pursner 2020). We highlight here some of the particularly important choices made while developing the framework (in order of appearance in the pipeline **Fig. 1**):

1. Steps for curation of eBird data (**1., Curating eBird data**) in addition to those outlined in Johnston et al. (2019), in particular to review checklist completeness.
2. Systematic objective process to remove vagrants from large datasets (**1., Reshaping eBird data, Removing vagrant records**), an important step to ensure that the data used for the analysis of any species can be constrained to be from within the expected distribution range of that species (**1., Selecting species and checklists, Constraining data for a species**). This is a vital step because the inclusion of data from a time period when a species is not present and from a region where it does not occur will make the estimated frequencies more susceptible to sampling biases over space and time. For a migratory species, not including this step will mean that checklists uploaded in the breeding region from the non-breeding season (when it cannot be present in the area) will also be used in the estimation of the species’ trends.
3. Spatial subsampling to reduce the spatial non-independence of checklists (**1., Selecting species and checklists, Reducing spatial non-independence of checklists**) in trend models. While SoIB (2020) only used random effects at the grid level to reduce spatial bias, it did not account for bias from variation in sampling between sites within each grid.
4. Specifying random effects as grids (**2., Modelling frequency of reporting, Running single-species models**) rather than hotspots/locations (e.g., Horns et al. 2018, Neate-Clegg et al. 2020, Kittelberger et al. 2023) in GLMMs. On eBird, hotspots and locations are often clustered together in the best sampled parts of a region, often around urban centres. Specifying random effects using checklist locations will therefore reduce the impact of any single site on the result (see previous point) but will not reduce urban bias across a larger region. Any derived trend may therefore be biased by how the distribution of hotspot/location clusters have changed over time.
5. Specifying ‘year’ as a fixed effect, but as a non-numerical factor (**2., Modelling frequency of reporting, Running single-species models**) rather than a continuous variable (e.g., Neate-Clegg et al. 2020, Kittelberger et al. 2023). Species trends over the long term can fluctuate or have opposing trajectories during different periods. Specifying year as a continuous variable forces a monotonically increasing or decreasing trend.
6. Specifying ‘season’ as a fixed effect (**2., Modelling frequency of reporting, Running single-species models**), a step not followed in the other similar analyses. Detectability of a species can vary across seasons, and trends can therefore be biased by changing patterns of seasonal reporting on eBird, unless season is explicitly controlled for. For example, let’s assume a certain species becomes less detectable during winter. If patterns of seasonal birding have changed over time from more summer birding in the past to more winter birding today, frequencies of reporting will inherently reduce, indicating a decline even when there is none.
7. Sensitivity checks to ensure that the robustness of the GLMER model results (**2., Modelling frequency of reporting, Model confidence checks**), the LTC categorization, and the CAT categorization (**2., Estimating LTC and CAT, Sensitivity checks**).
8. Determining a three-way priority classification for each species (**4.**), in alignment with other conventional regional State of Birds assessments (e.g., Burns et al. 2020).

The framework we present is detailed, but we believe that such detail is essential when dealing with a dataset that has so much inherent variability in sampling effort, observer experience, and degree of curation across space and time.

### Open habitat birds, insectivores, and raptors in global peril

Wild bird populations have declined considerably worldwide, with North America reporting the loss of three billion individual birds in the last fifty years (Rosenberg et al. 2019), and Europe the loss of over 500 million since 1980 (Burns et al. 2021). SoIB 2023 paints a similar picture in India, one of severe long-term decline, although the number of birds lost is as yet unknown. A common thread that emerges from the majority of global assessments is that some groups of birds are more threatened than others, in particular those that specialize entirely on grassland, scrubland, agriculture, or other open landscapes (BirdLife International 2022a). Grassland ecosystems are highly vulnerable in India due to the loss of habitat to development and plantations (see SoIB 2023b), and agricultural landscapes are rapidly changing in the country (Brandt et al. 2024). Several regional assessments now report significant declines of farmland/grassland birds in the long term: ∼40% in North America (North American Bird Conservation Initiative 2022), ∼50% in Europe (Burns et al. 2020), ∼40% in Australia (BirdLife Australia 2015b), and now ∼50% in India. In Europe, these declines have been attributed to agricultural intensification, largely from increased and unregulated pesticide and fertilizer use (Rigal et al. 2023).

Consequences of pesticide use are poorly understood in India but pesticides have been implicated in the decline of open habitat birds and insectivores also in the US (Li et al. 2020), and are widely thought to be the driving factor in global insect declines (Sánchez-Bayo and Wyckhuys 2019). Neonicotinoids, a family of pesticides, has recently been under scrutiny due to its documented toxicity towards insects (Pisa et al. 2021), in particular towards bees (Woodcock et al. 2016). This scrutiny has resulted in the ban of three neonicotinoids in the EU by the European Commission in 2018 (European Commission 2018). Pesticides, including neonicotinoids, currently have very little regulation in India and may be a reason behind the particularly severe declines of open habitat birds and insectivores. The relationship of pesticides and fertilizers with insect and bird declines in India requires urgent attention so that necessary steps can be taken quickly.

The plight of vultures in India is now well documented, alongside evidence that many documented threats still remain (see Threats section in SoIB 2023b). We worryingly found that most vulture species in India are in continued Rapid Decline in the country as a whole, with Indian Vulture showing the steepest declining trend. Prakash et al. (2024) reported that vulture trends have stabilized in many parts of the country. Results in SoIB differ, perhaps due to the inclusion of non-protected areas, and/or southern and northeastern India in this national average. Trends for vultures and raptors in general match trends from the African continent (Shaw et al. 2024), where all large vulture species have declined by more than 80% in the long term. In India and the African continent, Eurasian Kestrel, Tawny Eagle, and Montagu’s Harrier have rapidly declined (**Table 6**), while Booted Eagle has done well. Species that have declined in Africa but are faring reasonably well in India include Steppe Eagle, Black Kite, and Shikra. Western Marsh Harrier on the other hand is near-stable in Africa, but has declined rapidly in India (also see the results of systematic monitoring in Nannaj in Bharadwaj et al. (2024)).

**Table 6:** A comparison of the magnitudes of mean long-term change in the encounter rates/reporting frequency of similar raptor species in Africa (Shaw et al. 2024) and in India. When two species are mentioned in the column “Common Name” as “species 1/ species 2”, species 1 refers to the species in Africa and species 2 to the equivalent species in India.

| **Common Name** | **Africa:**<br>**LTC (%)** | **India:**<br>**LTC (%)** | **LTC Status** | **CAT Status** |
| --- | --- | --- | --- | --- |
| Steppe Eagle | -90 | -31 | Trend Inconclusive | Stable |
| White-backed Vulture/<br>White-rumped Vulture | -86 | -98 | Rapid Decline | Decline |
| Eurasian Kestrel | -70 | -72 | Rapid Decline | Rapid Decline |
| Tawny Eagle | -66 | -73 | Rapid Decline | Rapid Decline |
| Black Kite | -60 | 4 | Stable | Trend Inconclusive |
| Montagu's Harrier | -51 | -63 | Rapid Decline | Decline |
| Shikra | -44 | 22 | Trend Inconclusive | Stable |
| Black-winged Kite | -32 | -43 | Decline | Decline |
| Red-necked Falcon | -26 | -57 | Decline | Decline |
| Short-toed Snake-Eagle | -25 | -63 | Rapid Decline | Rapid Decline |
| Western Marsh Harrier | -4 | -61 | Rapid Decline | Decline |
| Booted Eagle | 3 | -11 | Stable | Trend Inconclusive |

### Contributions to global assessments

SoIB trends can play an important role in informing global assessments of resident bird species that are largely restricted to the region, or have sizeable populations within India. Declining trends for Western Ghats endemics indicate that their global statuses may require scrutiny. On the other hand, increasing trends in India may also have global implications. In the latest call for IUCN status reassessments, Birdlife International has referred to the increasing SoIB trend of Black-headed Ibis while proposing that its status be downlisted to Least Concern from Near Threatened (Red List Team (BirdLife International) 2024a).

India also harbours sizeable wintering populations of many species that breed in regions that are not well monitored. SoIB therefore provides insights not just into the status of resident birds within India, but also into the global status of many migratory species. An example is Black-capped Kingfisher, a winter migrant that had shown (and continues to show) evidence of rapid decline in India (SoIB 2020). Some evidence of decline from its breeding range in South Korea (Kim et al. 2021), combined with evidence from non-breeding regions including the SoIB trend, meant that the global IUCN status of the species was uplisted to Vulnerable (Red List Team (BirdLife International) 2022). SoIB 2023 provides insights into several other species whose wintering abundance is declining in India, but are globally under the radar such as Baillon’s Crake, Forest Wagtail, and Spot-winged Starling.

With its large coastline and network of wetlands, India is especially important to migratory shorebirds that use the Central Asian Flyway and the East Asian-Australasian Flyway. Our results add to the growing evidence that arctic breeding shorebirds are severely threatened, even in northern Russia where the birds that winter in India presumably breed. Of the 41 species of largely migratory shorebirds considered in SoIB, 25 (60%) are categorized as of High Priority. Arctic-breeding shorebirds (long-distance Arctic migrants) have declined by close to 80% as a group, considerably more than those that are near-resident or are Palearctic migrants (**Fig. 9**). In the latest call for IUCN status reassessments, Birdlife International has referred to the declining SoIB trends of Black-bellied Plover, Ruddy Turnstone, and Curlew Sandpiper while proposing that their status be uplisted (Red List Team (BirdLife International) 2024c, d, b).

**Fig. 9:**
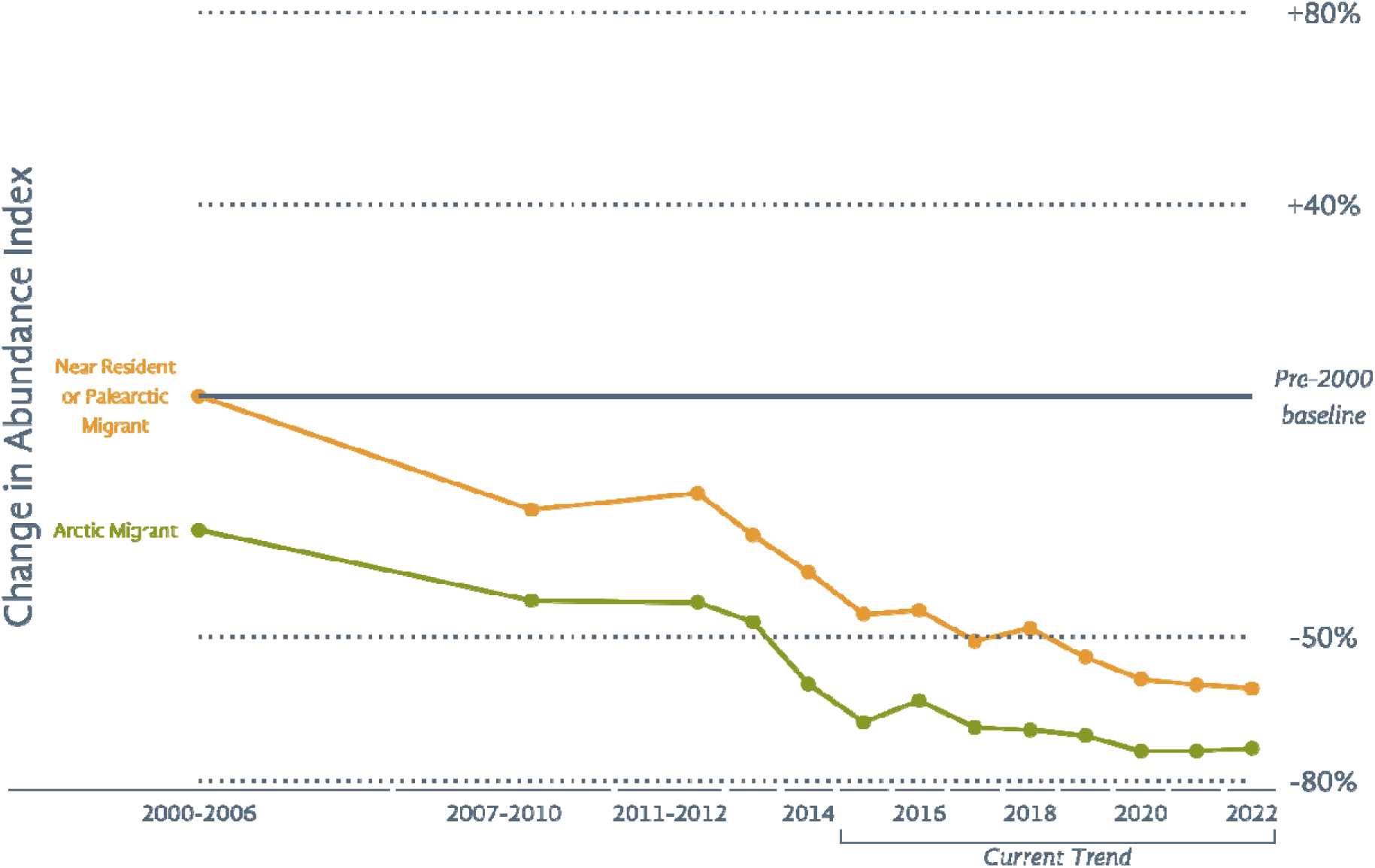
A comparison of the composite trends of shorebirds based on their migratory behaviours. Composite trends were calculated by averaging the mean individual trends of several species. Points refer to the proportional change in reporting frequency when compared to the pre-2000 baseline. Note that CIs have not been depicted. For detailed information about how to interpret this graph and for the lists of constituent species included in the composites, see the Methods section in the SoIB 2023 report (SoIB 2023b).

Populations of migratory ducks are in rapid decline in India, providing insights into source populations that breed east of Europe. Rates of decline appear very similar to those in Europe (**Table 7**). Duck species that have declined the least in SoIB like Green-winged Teal and Red-crested Pochard are the only ones showing increasing trends in Europe. Our results suggest that ducks and wetlands require focused conservation attention across Eurasia, much like the focused measures in North America that have enabled a remarkable recovery of waterfowl in the region (North American Bird Conservation Initiative 2022).

**Table 7:** A comparison of the magnitudes of mean long-term change in the population size/reporting frequency of ten duck species in Europe (Burns et al. 2021) and in India. All these species are winter migrants to the Indian subcontinent.

| Common Name | Europe:<br>LTC (%) | India:<br>LTC (%) | LTC Status | CAT Status |
| --- | --- | --- | --- | --- |
| Northern Pintail | -75 | -75 | Rapid Decline | Rapid Decline |
| Eurasian Wigeon | -60 | -60 | Decline | Decline |
| Garganey | -50 | -58 | Rapid Decline | Rapid Decline |
| Common Pochard | -50 | -68 | Rapid Decline | Decline |
| Northern Shoveler | -40 | -65 | Rapid Decline | Rapid Decline |
| Tufted Duck | -40 | -78 | Rapid Decline | Decline |
| Green-winged Teal | -25 | -69 | Rapid Decline | Rapid Decline |
| Red-crested Pochard | 25 | -42 | Decline | Stable |
| Gadwall | 50 | -27 | Trend Inconclusive | Decline |
| Ferruginous Duck |  | -40 | Trend Inconclusive | Trend Inconclusive |

### Systematic monitoring efforts remain essential

SoIB 2023 has limitations, largely due to the way eBird data is collected. We do not present estimates of population size because of the semi-structured nature of eBird data and non-standardized counting techniques. Several assessments around the world that are built on systematic monitoring efforts (Burns et al. 2020, North American Bird Conservation Initiative 2022) report population sizes adding an important dimension in the prioritization of bird species for conservation. We encourage the initiation of long-term systematic efforts in India that can supplement eBird data and address this gap. Some assessments like the State of the UK’s Birds (Burns et al. 2020) also report trends in demographic parameters, an important metric that can give early warning signals of decline especially for long-lived birds like vultures and storks. Demographic trends, however, cannot be derived from eBird data, and require dedicated long-term systematic monitoring. At least one long-term study in India monitors demography (Srinivasan and Wilcove 2021), and this can hopefully be replicated and scaled up in the future to produce the required data at the national level.

SoIB currently lacks abundance trends for certain groups of birds like mountain pheasants, nocturnal birds (owls, nightjars), and seabirds. These groups are poorly represented in citizen science (or other) datasets for large scale analysis, often because of low detectability. Specialised networks focussed on monitoring these groups are necessary to understand their trends at a national scale, as is the case in the UK (Burns et al. 2020) where even the Breeding Bird Survey is insufficient to monitor rare birds. SoIB also has very few available results for range restricted species (**Table 3**), due to a lack of data as well as limitations in the analytical methodology that make the analyses of restricted species difficult. Data uploaded to eBird is collected by birders with a wide range of experience, leading to poor data for species where identification is especially challenging. Many such species are not considered for LTC and CAT analysis in SoIB, but some estimated frequencies for bird groups like pipits and *Acrocephalus*/*Iduna* warblers may need to be treated with caution. Systematic and extensive ringing studies may be essential for estimating abundances of difficult groups of birds such as these.

### Recommendations for citizen science and conservation practitioners in emerging regions

Birdwatching is growing in popularity worldwide, creating an opportunity for citizen science projects to blossom in regions where baseline data on birds are lacking. While the step from casual birdwatching to more formal citizen science can be organic, we urge concerned people in the community to become citizen science managers and accelerate the growth of citizen science through dedicated programmes and workshops. Such an investment ensures significant gains both in terms of monitoring process and conservation outcome. Deepening citizen participation in an open and transparent monitoring of biodiversity at massive scales is highly desirable in and of itself. But in addition, such an effort yields the data necessary to produce assessments of the State of Birds, especially when coupled with the framework we have put forward to enhance their rigour and reliability.

Institutionally too, the State of India’s Birds 2023 report (SoIB 2023b) offers a collaborative model whereby thirteen governmental and non-governmental organizations, each with their particular expertise have partnered around a common set of objectives and approaches. We recommend a similar approach for a State of Birds report elsewhere too, because it results in an assessment that is not only scientifically reliable, but also institutionally robust, offering greater legitimacy for its results to be integrated into downstream public decision-making for conservation.

## Acknowledgements

We thank the entire community of eBirders and data quality editors whose contributions have made this assessment possible. Certificates were sent to all contributors highlighting their efforts. The authors would like to thank the remaining members of the partnership who were also involved in the making of the report: Ananda Banerjee, Anup Bokkasa, Aparajita Datta, Arghya Chakrabarty, B Balaji, Biswajit Chakdar, Bivash Pandav, Dhriti Banerjee, Dhruv Verma, Dipankar Ghose, Diwakar Sharma, Farah Ishtiaq, Gautam Talukdar, Gopi GV, Gopinathan Maheswaran, Himani Sharma, Janhavi Rajan, K Ramesh, Kiran Singh, LS Shashidhara, Madhumita Panigrahi, Malvika Onial, Merwyn Fernandes, Neha Sinha, Parveen Shaikh, PO Nameer, Prachi Mehta, Rahul Kaul, Rahul Kishor Talegaonkar, Rajkamal Goswami, Raju Kasambe, Ramesh Kumar Selvaraj, Ravi Singh, Ritesh Kumar, Rohit Jha, S Sathyakumar, Sejal Worah, Sanjay Molur, Subhadra Devi, Sujit Narwade, Suresh Kumar, Taej Mundkur, Uma Ramakrishnan, Umesh Srinivasan, Vijay Ramesh, Vishnupriya Kolipakam, Vinay KL, and Vivek Menon. We are also grateful to Alex Berryman, Alison Johnston, Asad Rahmani, Rob Martin, Subbu Subramanya, TR Shankar Raman, Wesley M Hochachka, and Yong Ding Li for their critical comments and technical inputs. We gratefully acknowledge financial support from Duleep Matthai Nature Conservation Trust, Foundation for Ecological Security, Rohini Nilekani Philanthropies, The Habitats Trust, Wildlife Trust of India, World Wide Fund for Nature–India.

## References

Ali, S., and S. D. Ripley. 1968. Handbook of the Birds of India and Pakistan: Together with Those of Nepal, Sikkim, Bhutan and Ceylon 10V. Oxford University Press.

Altwegg, R., and J. D. Nichols. 2019. Occupancy models for citizen-science data. Methods in Ecology and Evolution 10:8–21.

Barrett, G., A. Silcocks, S. Barry, R. Cunningham, and R. Poulter. 2003. The new atlas of Australian birds. Royal Australasian Ornithologists Union, Melbourne.

Bates, D., M. Maechler, B. Bolker, and S. Walker. 2015. Fitting Linear Mixed-Effects Models Using lme4. Journal of Statistical Software 67:1–48.

Bharadwaj, A., S. Mhamane, P. Bangal, T. Menon, K. Isvaran, and S. Quader. 2024. Assessment of long-term trends in a threatened grassland bird community using daily bird lists. Bird Conservation International 34:e3.

Bird, T. J., A. E. Bates, J. S. Lefcheck, N. A. Hill, R. J. Thomson, G. J. Edgar, R. D. Stuart-Smith, S. Wotherspoon, M. Krkosek, J. F. Stuart-Smith, G. T. Pecl, N. Barrett, and S. Frusher. 2014. Statistical solutions for error and bias in global citizen science datasets. Biological Conservation 173:144–154.

BirdLife Australia. 2015a. Measuring the state of Australia’s terrestrial birds: how Australian Bird Indices (ABIs) are made. Carlton, Australia.

BirdLife Australia. 2015b. The state of Australia’s birds 2015: headline trends for terrestrial birds. Carlton, Australia.

BirdLife International. 2015. European Red List of Birds. Luxembourg: Office for Official Publications of the European Communities.

BirdLife International. 2017. European birds of conservation concern: populations, trends and national responsibilities. Cambridge, UK.

BirdLife International. 2018. State of the world’s birds: taking the pulse of the planet. Cambridge, UK.

BirdLife International. 2022a. State of the World’s Birds 2022: Insights and Solutions for the Biodiversity Crisis. BirdLife International Cambridge, UK.

BirdLife International. 2022b. State of the World’s Birds: Insights and solutions for the biodiversity crisis. Cambridge, UK.

Blancher, P. J., R. D. Phoenix, D. S. Badzinski, M. D. Cadman, T. L. Crewe, C. M. Downes, D. Fillman, C. M. Francis, J. Hughes, D. J. T. Hussell, D. Lepage, J. D. McCracken, D. K. McNicol, B. A. Pond, R. K. Ross, R. Russell, L. A. Venier, and R. C. Weeber. 2009. Population trend status of Ontario’s forest birds. The Forestry Chronicle 85:184–201.

Brandt, M., D. Gominski, F. Reiner, A. Kariryaa, V. B. Guthula, P. Ciais, X. Tong, W. Zhang, D. Govindarajulu, D. Ortiz-Gonzalo, and R. Fensholt. 2024. Severe decline in large farmland trees in India over the past decade. Nature Sustainability.

British Trust for Ornithology, 2019. Webpage URL: https://www.bto.org/volunteer-surveys/bbs/research-conservation/methodology. [Accessed on 16 April 2024].

Brotons, L., and S. Herrando. 2011. Population Estimates: Towards Standardised Protocols as a Basis for Comparability. BIOONE.

Buckland, S. T., A. E. Magurran, R. E. Green, and R. M. Fewster. 2005. Monitoring change in biodiversity through composite indices. Philosophical Transactions of the Royal Society B: Biological Sciences 360:243–254.

Burns, F., M. A. Eaton, D. E. Balmer, A. Banks, R. Caldow, J. L. Donelan, A. Douse, C. Duigan, S. Foster, T. Frost, P. V. Grice, C. Hall, H. J. Hanmer, S. J. Harris, I. Johnstone, P. Lindley, N. McCulloch, D. G. Noble, K. Risely, R. A. Robinson, and S. Wotton. 2020. The state of the UK’s birds 2020. The RSPB, BTO, WWT, DAERA, JNCC, NatureScot, NE and NRW, Sandy, Bedfordshire.

Burns, F., M. A. Eaton, I. J. Burfield, A. Klvaňová, E. Šilarová, A. Staneva, and R. D. Gregory. 2021. Abundance decline in the avifauna of the European Union reveals cross-continental similarities in biodiversity change. Ecology and Evolution 11:16647–16660.

Clements, J. F., P. C. Rasmussen, T. S. Schulenberg, M. J. Iliff, T. A. Fredericks, J. A. Gerbracht, D. Lepage, A. Spencer, S. M. Billerman, B. L. Sullivan, and C. L. Wood. 2022. The eBird/Clements checklist of birds of the world: v2022. Downloaded from http://www.birds.cornell.edu/clementschecklist/download/.

Clements, J. F., P. C. Rasmussen, T. S. Schulenberg, M. J. Iliff, T. A. Fredericks, J. A. Gerbracht, D. Lepage, A. Spencer, S. M. Billerman, B. L. Sullivan, and C. L. Wood. 2023. The eBird/Clements checklist of birds of the world: v2023. Downloaded from http://www.birds.cornell.edu/clementschecklist/download/.

Crowe, O., A. J. Musgrove, and J. O’Halloran. 2014. Generating population estimates for common and widespread breeding birds in Ireland. Bird Study 61:82–90.

Díaz, S., J. Settele, E. Brondízio, H. T. Ngo, M. Guèze, J. Agard, A. Arneth, P. Balvanera, K. Brauman, S. Butchart, K. Chan, L. Garibaldi, K. Ichii, J. Liu, S. M. Subramanian, G. Midgley, P. Miloslavich, Z. Molnár, D. Obura, A. Pfaff, S. Polasky, A. Purvis, J. Razzaque, B. Reyers, R. R. Chowdhury, Y.-J. Shin, I. Visseren-Hamakers, K. Willis, and C. Zayas. 2019. Summary for policymakers of the global assessment report on biodiversity and ecosystem services of the Intergovernmental Science-Policy Platform on Biodiversity and Ecosystem Services. Intergovernmental Science-Policy Platform on Biodiversity and Ecosystem Services (IPBES), Bonn, Germany.

Eaton, M., N. Aebischer, A. Brown, R. Hearn, L. Lock, A. Musgrove, D. Noble, D. Stroud, and R. Gregory. 2015a. Birds of Conservation Concern 4: the population status of birds in the UK, Channel Islands and Isle of Man. British Birds 108:708–746.

Eaton, M., N. Aebischer, A. Brown, R. Hearn, L. Lock, A. Musgrove, D. Noble, D. Stroud, and R. Gregory. 2015b. Supplementary Material - Birds of Conservation Concern 4: the population status of birds in the UK, Channel Islands and Isle of Man. British Birds 108:708–746. eBird, 2024. Webpage URL: https://support.ebird.org/en/support/solutions/articles/48000950859-guide-to-ebird-protocols. [Accessed on 16 April 2024].

European Bird Census Council (EBCC), 2018. Webpage URL: http://www.ebcc.info/methods-2018/. [Accessed on 16 April 2024].

European Commission, 2018. Webpage URL: https://food.ec.europa.eu/plants/pesticides/approval-active-substances/renewal-approval/neonicotinoids_en.

Fiske, I., and R. Chandler. 2011. unmarked: An R Package for Fitting Hierarchical Models of Wildlife Occurrence and Abundance. 2011 43:23.

Gregory, R. D., A. van Strien, P. Vorisek, W. Gmelig Meyling Adriaan, G. Noble David, P. B. Foppen Ruud, and W. Gibbons David. 2005. Developing indicators for European birds. Philosophical Transactions of the Royal Society B: Biological Sciences 360:269–288.

Grimmett, R., C. Inskipp, and T. Inskipp. 2011. Birds of the Indian Subcontinent. Oxford University Press.

Harris, S. J., D. Massimino, S. Gillings, M. A. Eaton, D. G. Noble, D. E. Balmer, D. Procter, P.-H. J.W., and P. Woodcock. 2018. The Breeding Bird Survey 2017. Thetford.

Hayhow, D. B., M. A. Ausden, R. B. Bradbury, D. Burnell, A. I. Copeland, H. Q. P. Crick, M. A. Eaton, T. Frost, P. V. Grice, C. Hall, S. J. Harris, M. D. Morecroft, D. G. Noble, P.-H. J.W., O. Watts, and J. M. Williams. 2017. The state of the UK’s birds 2017. The RSPB, BTO, WWT, DAERA, JNCC, NE and NRW, Sandy, Bedfordshire.

Herrando, S., L. Brotons, J. Estrada, and V. Pedrocchi. 2008. The Catalan Common Bird Survey (SOCC): a tool to estimate species population numbers.

Horns, J. J., F. R. Adler, and Ç. H. Şekercioğlu. 2018. Using opportunistic citizen science data to estimate avian population trends. Biological Conservation 221:151–159.

IUCN. 2023. The IUCN Red List of Threatened Species. Version 2023-1. http://www.iucnredlist.org. Accessed on 20 March 2023.

Johnston, A., D. Fink, W. M. Hochachka, and S. Kelling. 2018. Estimates of observer expertise improve species distributions from citizen science data. Methods in Ecology and Evolution 9:88–97.

Johnston, A., W. M. Hochachka, M. Strimas-Mackey, V. Ruiz Gutierrez, O. J. Robinson, E. T. Miller, T. Auer, S. T. Kelling, and D. Fink. 2019. Best practices for making reliable inferences from citizen science data: case study using eBird to estimate species distributions. bioRxiv:574392.

Joseph, L. N., S. A. Field, C. Wilcox, and H. P. possingham. 2006. Presence–Absence versus Abundance Data for Monitoring Threatened Species. Conservation Biology 20:1679–1687.

Kelling, S., A. Johnston, W. M. Hochachka, M. Iliff, D. Fink, J. Gerbracht, C. Lagoze, F. A. La Sorte, T. Moore, A. Wiggins, W.-K. Wong, C. Wood, and J. Yu. 2015. Can Observation Skills of Citizen Scientists Be Estimated Using Species Accumulation Curves? PLoS ONE 10:e0139600.

Kéry, M., B. Gardner, and C. Monnerat. 2010. Predicting species distributions from checklist data using site-occupancy models. Journal of Biogeography 37:1851–1862.

Kim, H., Y. Mo, C.-Y. Choi, B. C. McComb, and M. G. Betts. 2021. Declines in Common and Migratory Breeding Landbird Species in South Korea Over the Past Two Decades. Frontiers in Ecology and Evolution 9.

Kittelberger, K. D., C. J. Tanner, N. D. Orton, and Ç. H. Şekercioğlu. 2023. The value of community science data to analyze long-term avian trends in understudied regions: The state of birds in Türkiye. Avian Research 14:100140.

Knaus, P., S. Antoniazza, S. Wechsler, J. Guélat, M. Kéry, N. Strebel, and T. Sattler. 2018. Swiss Breeding Bird Atlas 2013–2016. Distribution and population trends of birds in Switzerland and Liechtenstein. Swiss Ornithological Institute, Sempach.

Knowles, J. E., C. Frederick, A. Whitworth, and M. J. E. Knowles. 2023. Package ‘merTools’.

Lees, A. C., L. Haskell, T. Allinson, S. B. Bezeng, I. J. Burfield, L. M. Renjifo, K. V. Rosenberg, A. Viswanathan, and S. H. M. Butchart. 2022. State of the World’s Birds. Annual Review of Environment and Resources 47:231–260.

Li, Y., R. Miao, and M. Khanna. 2020. Neonicotinoids and decline in bird biodiversity in the United States. Nature Sustainability 3:1027–1035.

Lin, D.-l., and S. Pursner. 2020. The State of Taiwan’s Birds 2020.

Link, W. A., and J. R. Sauer. 1998. Estimating population change from count data: Application to the north american breeding bird survey. Ecological Applications 8:258–268.

Magurran, A. E., and P. A. Henderson. 2003. Explaining the excess of rare species in natural species abundance distributions. Nature 422:714–716.

Michigan State University, 2019. Webpage URL: http://www.lon-capa.org/∼mmp/labs/error/e2.htm. [Accessed on 16 April 2024].

Moosmann, M., N. Auchli, T. Kuzmenko, T. Sattler, H. Schmid, B. Volet, S. Wechsler, and N. Strebel. 2023. The State of Birds in Switzerland: Report 2023. Swiss Ornithological Institute, Sempach.

Musgrove, A., N. Aebischer, M. Eaton, R. Hearn, S. Newson, D. Noble, M. Parsons, K. Risely, and D. Stroud. 2013. Population estimates of birds in Great Britain and the United Kingdom.

Neate-Clegg, M. H. C., J. J. Horns, F. R. Adler, M. Ç. Kemahlı Aytekin, and Ç. H. Şekercioğlu. 2020. Monitoring the world’s bird populations with community science data. Biological Conservation 248:108653.

Newson, S. E., K. L. Evans, D. G. Noble, J. J. D. Greenwood, and K. J. Gaston. 2008. Use of distance sampling to improve estimates of national population sizes for common and widespread breeding birds in the UK. Journal of Applied Ecology 45:1330–1338.

Newson, S. E., R. J. W. Woodburn, D. G. Noble, S. R. Baillie, and R. D. Gregory. 2005. Evaluating the Breeding Bird Survey for producing national population size and density estimates. Bird Study 52:42–54.

Norman, D., R. J. Harris, and S. E. Newson. 2012. Producing regional estimates of population size for common and widespread breeding birds from national monitoring data. Bird Study 59:10–21.

North American Bird Conservation Initiative. 2012. The state of Canada’s birds, 2012. Ottawa, Canada.

North American Bird Conservation Initiative. 2022. The State of the Birds, United States of America, 2022. U.S. Department of Interior, Washington, D.C.

North American Bird Conservation Initiative, U. S. C. 2014. The State of the Birds 2014 Report. U.S. Department of Interior, Washington, D.C.

Olsen, P., M. Weston, and A. Silcocks. 2003. The State of Australia’s Birds 2003.

Pisa, L., D. Goulson, E.-C. Yang, D. Gibbons, F. Sánchez-Bayo, E. Mitchell, A. Aebi, J. van der Sluijs, C. J. K. MacQuarrie, C. Giorio, E. Y. Long, M. McField, M. Bijleveld van Lexmond, and J.-M. Bonmatin. 2021. An update of the Worldwide Integrated Assessment (WIA) on systemic insecticides. Part 2: impacts on organisms and ecosystems. Environmental Science and Pollution Research 28:11749–11797.

Pollock, J. F. 2006. Detecting Population Declines over Large Areas with Presence-Absence, Time-to-Encounter, and Count Survey Methods. Conservation Biology 20:882–892.

Prakash, V., H. Bajpai, S. S. Chakraborty, M. Singh Mahadev, J. W. Mallord, N. Prakash, S. P. Ranade, R. N. Shringarpure, C. G. R. Bowden, and R. E. Green. 2024. Recent trends in populations of Critically Endangered Gyps vultures in India. Bird Conservation International 34:e1.

Praveen, J., and R. Jayapal. 2023. Checklist of the birds of India (v7.1). Website: http://www.indianbirds.in/india/ [Date of publication: 28 February 2023].

R Core Team. 2023. R: A language and environment for statistical computing. R Foundation for Statistical Computing, Vienna, Austria.

Rasmussen, P. C., and J. C. Anderton. 2012. Birds of South Asia: The Ripley Guide: Attributes and Status. 2 edition. Smithsonian Institution and Lynx Edicions, Washington DC and Barcelona.

Red List Team (BirdLife International), 2022. Webpage URL: https://forums.birdlife.org/2022-1-black-capped-kingfisher-halcyon-pileata-revise-global-status/.

Red List Team (BirdLife International), 2024a. Webpage URL: https://forums.birdlife.org/2024-1-black-headed-ibis-threskiornis-melanocephalus/.

Red List Team (BirdLife International), 2024b. Webpage URL: https://forums.birdlife.org/2024-2-curlew-sandpiper-calidris-ferruginea/.

Red List Team (BirdLife International), 2024c. Webpage URL: https://forums.birdlife.org/2024-2-grey-plover-pluvialis-squatarola/.

Red List Team (BirdLife International), 2024d. Webpage URL: https://forums.birdlife.org/2024-2-ruddy-turnstone-arenaria-interpres/.

Rigal, S., V. Dakos, H. Alonso, A. Auniņš, Z. Benkő, L. Brotons, T. Chodkiewicz, P. Chylarecki, E. de Carli, J. C. del Moral, C. Domşa, V. Escandell, B. Fontaine, R. Foppen, R. Gregory, S. Harris, S. Herrando, M. Husby, C. Ieronymidou, F. Jiguet, J. Kennedy, A. Klvaňová, P. Kmecl, L. Kuczyński, P. Kurlavičius, J. A. Kålås, A. Lehikoinen, Å. Lindström, R. Lorrillière, C. Moshøj, R. Nellis, D. Noble, D. P. Eskildsen, J.-Y. Paquet, M. Pélissié, C. Pladevall, D. Portolou, J. Reif, H. Schmid, B. Seaman, Z. D. Szabo, T. Szép, G. T. Florenzano, N. Teufelbauer, S. Trautmann, C. van Turnhout, Z. Vermouzek, T. Vikstrøm, P. Voříšek, A. Weiserbs, and V. Devictor. 2023. Farmland practices are driving bird population decline across Europe. Proceedings of the National Academy of Sciences 120:e2216573120.

Rosenberg, K. V., A. M. Dokter, P. J. Blancher, J. R. Sauer, A. C. Smith, P. A. Smith, J. C. Stanton, A. Panjabi, L. Helft, M. Parr, and P. P. Marra. 2019. Decline of the North American avifauna. Science 366:120–124.

Sánchez-Bayo, F., and K. A. G. Wyckhuys. 2019. Worldwide decline of the entomofauna: A review of its drivers. Biological Conservation 232:8–27.

Sattler, T., P. Knaus, H. Schmid, and B. Volet. 2017. The State of Birds in Switzerland Report 2017. Swiss Ornithological Institute, Sempach.

Sauer, J. R., W. A. Link, J. E. Fallon, K. L. Pardieck, and J. David J. Ziolkowski. 2013. The North American Breeding Bird Survey 1966–2011: Summary Analysis and Species Accounts. North American Fauna:1–32.

Sauer, J. R., W. A. Link, and B. G. Peterjohn. 1994. Observer Differences in the North American Breeding Bird Survey. The Auk: Ornithological Advances 111:50–62.

Shaw, P., D. Ogada, L. Dunn, R. Buij, A. Amar, R. Garbett, M. Herremans, M. Z. Virani, C. J. Kendall, B. M. Croes, M. Odino, S. Kapila, P. Wairasho, C. Rutz, A. Botha, U. Gallo-Orsi, C. Murn, G. Maude, and S. Thomsett. 2024. African savanna raptors show evidence of widespread population collapse and a growing dependence on protected areas. Nature Ecology & Evolution 8:45–56.

SoIB. 2020. State of India’s Birds, 2020: Range, trends and conservation status. Pp. 50. The SoIB Partnership.

SoIB, 2023a. Webpage URL: https://stateofindiasbirds.in/. [Accessed on 16 April 2024].

SoIB. 2023b. State of India’s Birds, 2023: Range, trends, and conservation status. Pp. 119. The SoIB Partnership. doi:10.5281/zenodo.11124590.

Srinivasan, U., and D. S. Wilcove. 2021. Interactive impacts of climate change and land-use change on the demography of montane birds. Ecology 102:e03223.

stateofindiasbirds, 2024. Webpage URL: https://github.com/stateofindiasbirds/soib_2023. [Accessed on 16 April 2024].

Sullivan, B. L., J. L. Aycrigg, J. H. Barry, R. E. Bonney, N. Bruns, C. B. Cooper, T. Damoulas, A. A. Dhondt, T. Dietterich, A. Farnsworth, D. Fink, J. W. Fitzpatrick, T. Fredericks, J. Gerbracht, C. Gomes, W. M. Hochachka, M. J. Iliff, C. Lagoze, F. A. La Sorte, M. Merrifield, W. Morris, T. B. Phillips, M. Reynolds, A. D. Rodewald, K. V. Rosenberg, N. M. Trautmann, A. Wiggins, D. W. Winkler, W.-K. Wong, C. L. Wood, J. Yu, and S. Kelling. 2014. The eBird enterprise: An integrated approach to development and application of citizen science. Biological Conservation 169:31–40.

Szabo, J. K., P. A. Vesk, P. W. J. Baxter, and H. P. Possingham. 2010. Regional avian species declines estimated from volunteer-collected long-term data using List Length Analysis. Ecological Applications 20:2157–2169.

Walker, J., and P. D. Taylor. 2017. Using eBird data to model population change of migratory bird species. Avian Conservation and Ecology 12.

Wikipedia contributors, 2024. Webpage URL: https://en.wikipedia.org/w/index.php?title=Water_year&oldid=1199128054. [Accessed on 11 April 2024].

Wilman, H., J. Belmaker, J. Simpson, C. de la Rosa, M. M. Rivadeneira, and W. Jetz. 2014. EltonTraits 1.0: Species-level foraging attributes of the world’s birds and mammals. Ecology 95:2027–2027.

Woodcock, B. A., N. J. B. Isaac, J. M. Bullock, D. B. Roy, D. G. Garthwaite, A. Crowe, and R. F. Pywell. 2016. Impacts of neonicotinoid use on long-term population changes in wild bees in England. Nature Communications 7:12459.

