## Supplementary material for "State of India’s Birds 2023: A framework to leverage semi-structured citizen science for bird conservation"

***Components of SoIB 2023 whose methods are not presented in the article***

Some components of the report on ‘Systematic monitoring’ and ’Major threats to birds in India’ use different primary bird datasets. Brief analytical methods for these sections are included in the report (SoIB 2023) but are not presented in this article. One more component that focuses on recommendations to the global IUCN Red List does use eBird data (specifically the current annual trend), but the methodology is not described here and will be separately published.

***Placing data in broad regions***

For countrywide analyses of an ecologically diverse country like India, it was important to ensure that the various ecological regions of the country are well represented in the data. We therefore categorized every observation into one of eight ecological regions by broadly classifying each district (county in eBird data) in the country into one of these regions. Districts were physically assigned to a region by examining satellite view on Google Maps, and then hierarchically classified according to **Table S2**. Because this classification was only intended to enable diagnostic examination of results across different regions in the country rather than draw inferences about these different regions, we used the simplest possible criteria.

***Relationship between reporting frequency and list length***

Population trend indices are prone to bias if certain aspects of birder behaviour have changed over time, like the proportions of various habitats covered, or median list length, or mean birder abilities. We cannot easily examine issues like changes in the proportion of different habitats covered (given that checklists often cover multiple habitats), or changes in birder ability (difficult to assess), but we do correct for changes in effort over time in the analyses. One aspect that we have not considered in these analyses is the possibility that the relationship between list length and detectability may vary in space. We explored these relationships for some bird species in some broad ecological regions. We found that the relationship between reporting frequency and list length was similar across 3 well-birded and significant ecological regions in the country, the coast, plains, and the Western Ghats (**Fig. S7**). We therefore decided not to include this complexity in our analyses (to avoid overparameterization), but we recommend considering its inclusion in any analyses that are more data-rich, especially for species that span many ecological zones in their distributions.

***Decisions take about specific species***

Nocturnal species were excluded from the study but Jerdon's Courser, though completely nocturnal, was included here owing to its conservation significance in the region. We also did not consider Besra, Horsfield’s Bushlark (Singing Bushlark), Common Flameback, Eastern Orphean Warbler, Richard’s Pipit, and Asian Palm Swift for LTC and CAT analyses due to persisting issues with identification.

***Constraints for categorization of trends***

A potential concern with our criteria for categorizing trends stems from the use of the ratio of the estimate for each year-interval over the base year-interval. Because the constituent numbers are frequencies, ranging from 0 to 1, large values of the ratio (e.g., >2, corresponding to >100% increase) are only mathematically possible when the denominator (base frequency) is small (e.g., less than 0.5). To check whether >100% increases were mathematically possible, we examined the base frequencies for all species selected for LCT and CAT analyses (N = 523, 643). In all cases they were below 0.5, approaching a maximum of 0.38 (**Fig. S9**), illustrating that large values of the ratio are not mathematically prohibited by our analysis.

***Model diagnostics***

We ran standard model validation checks for the GLMs and GLMMs and ensured that, for a random subset of 50 species, standardized residuals for the best model were roughly evenly distributed around zero and not particularly skewed in either direction. We also checked the linearity of the quantile-quantile plot for the same random subset of fits. Random effects were approximately normally distributed and had non-zero variances (>10^-4^).

***Mapping to other lists***

We maintained a sheet that mapped to the India checklist and Birdlife International taxonomies (Praveen and Jayapal 2023), IUCN Red List status (IUCN 2022), Indian Wildlife (Protection) Act (1972) schedules, CITES and CMS appendices (adapted from Praveen and Jayapal 2023) and one percent estimates of global waterbird populations (Wetlands International 2012).

**Supplementary figures**
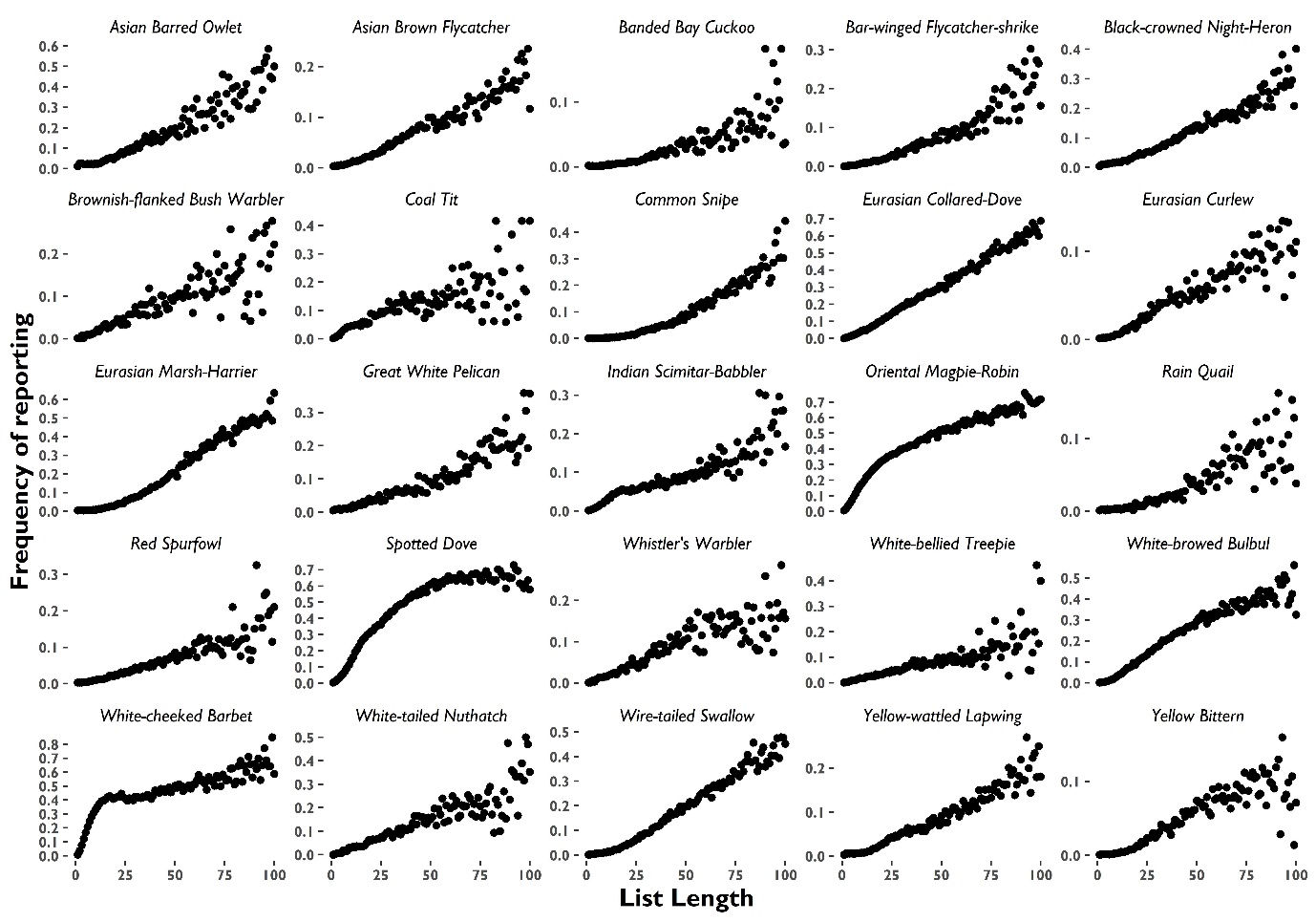

**Fig. S1**: Frequency of reporting of the first 25 of 50 randomly selected species in relation to list length (number of species in a checklist up to 100).

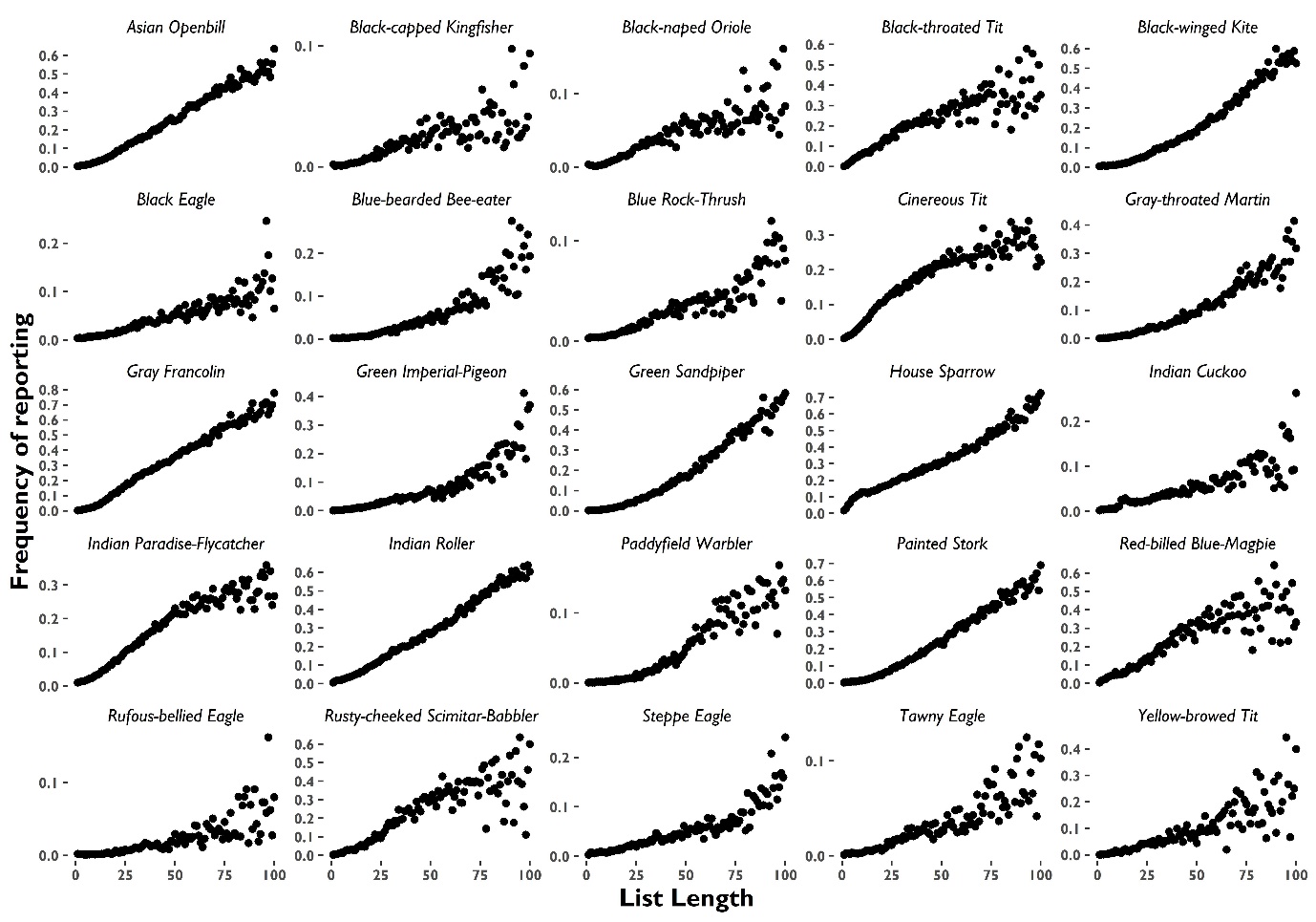

**Fig. S2**: Frequency of reporting of the second 25 of 50 randomly selected species in relation to list length (number of species in a checklist up to 100).

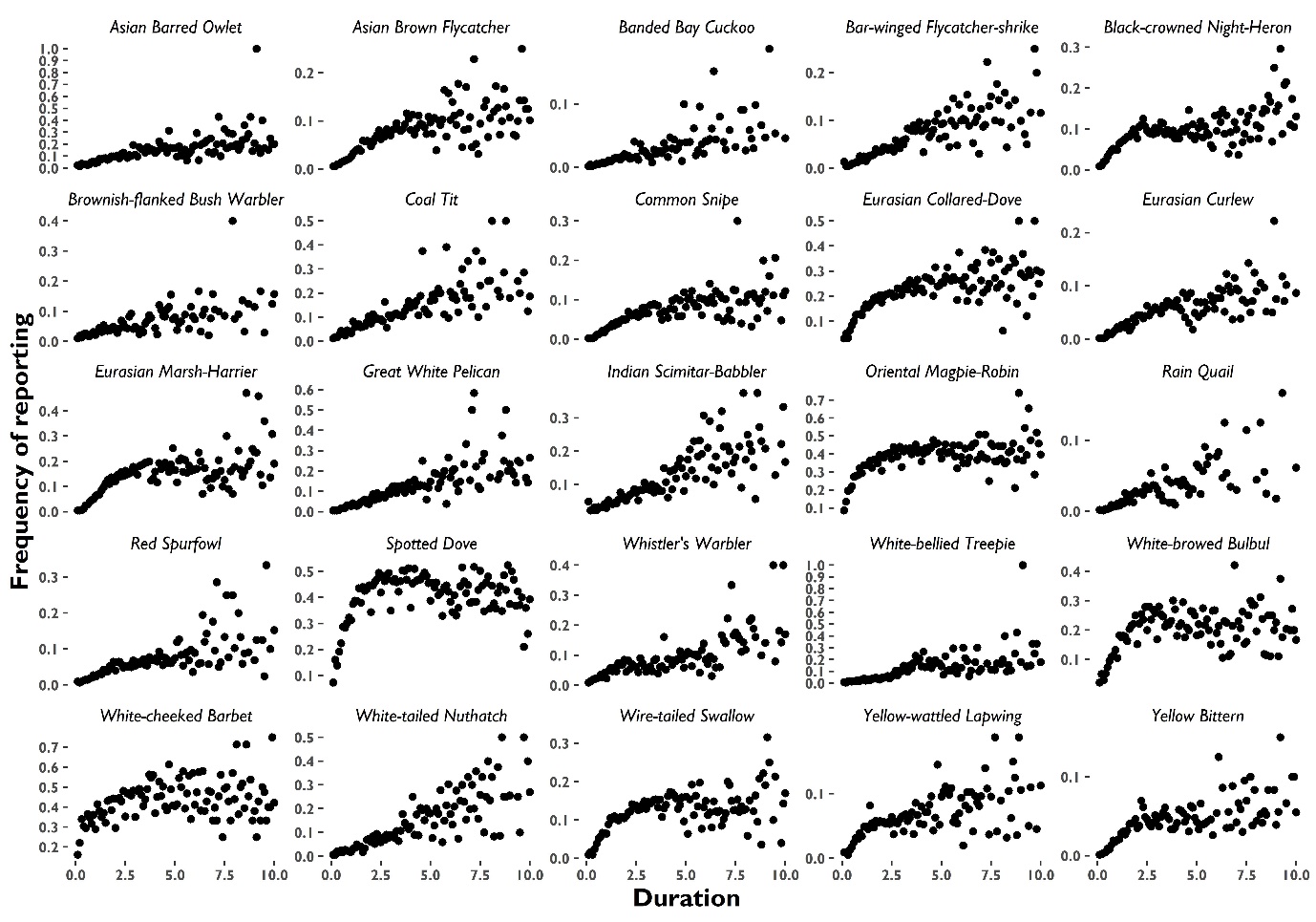

**Fig. S3**: Frequency of reporting of the first 25 of 50 randomly selected species in relation to binned Duration (100 0.1 hrs bins).

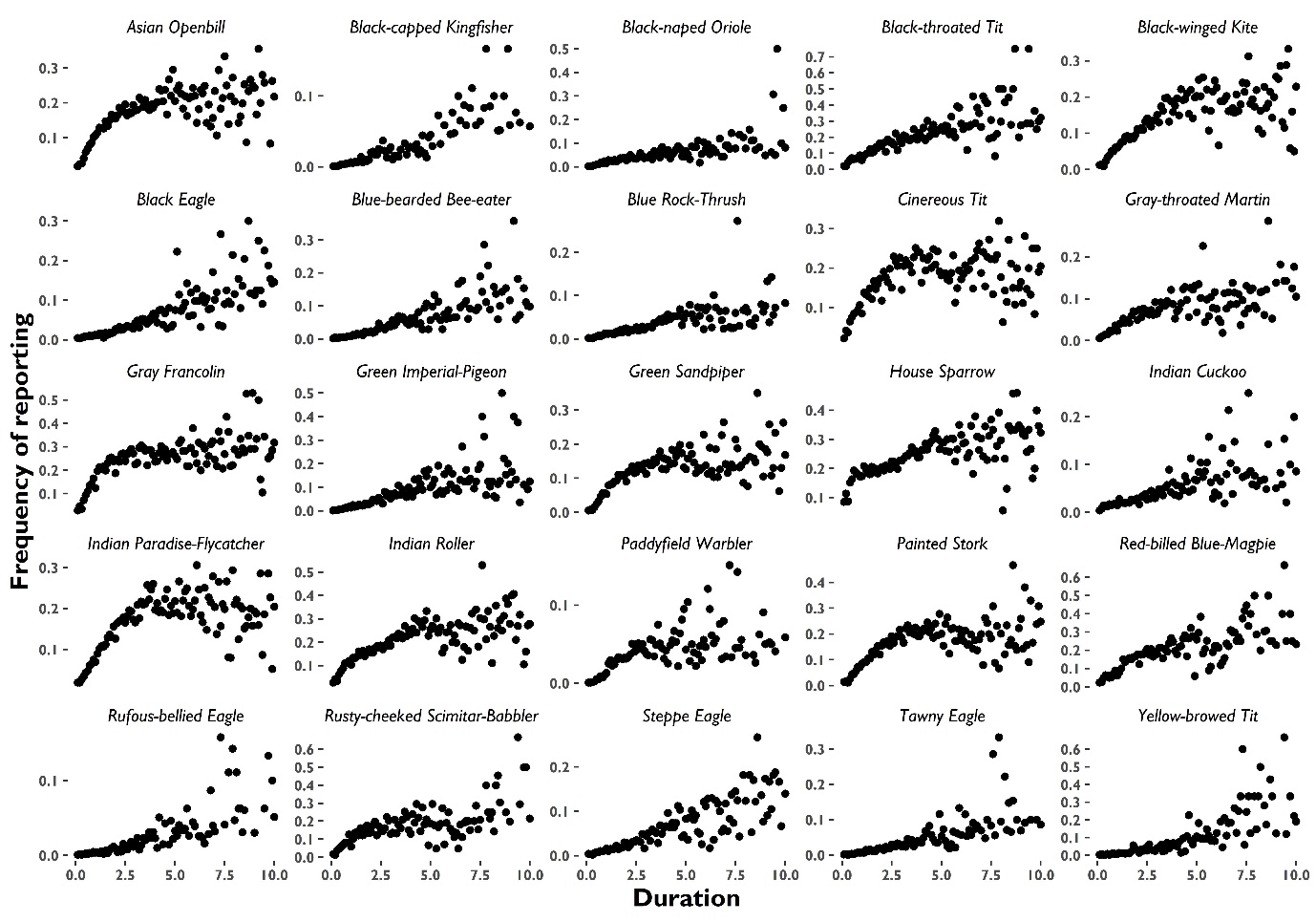

**Fig. S4**: Frequency of reporting of the second 25 of 50 randomly selected species in relation to binned Duration (100 0.1 hrs bins).

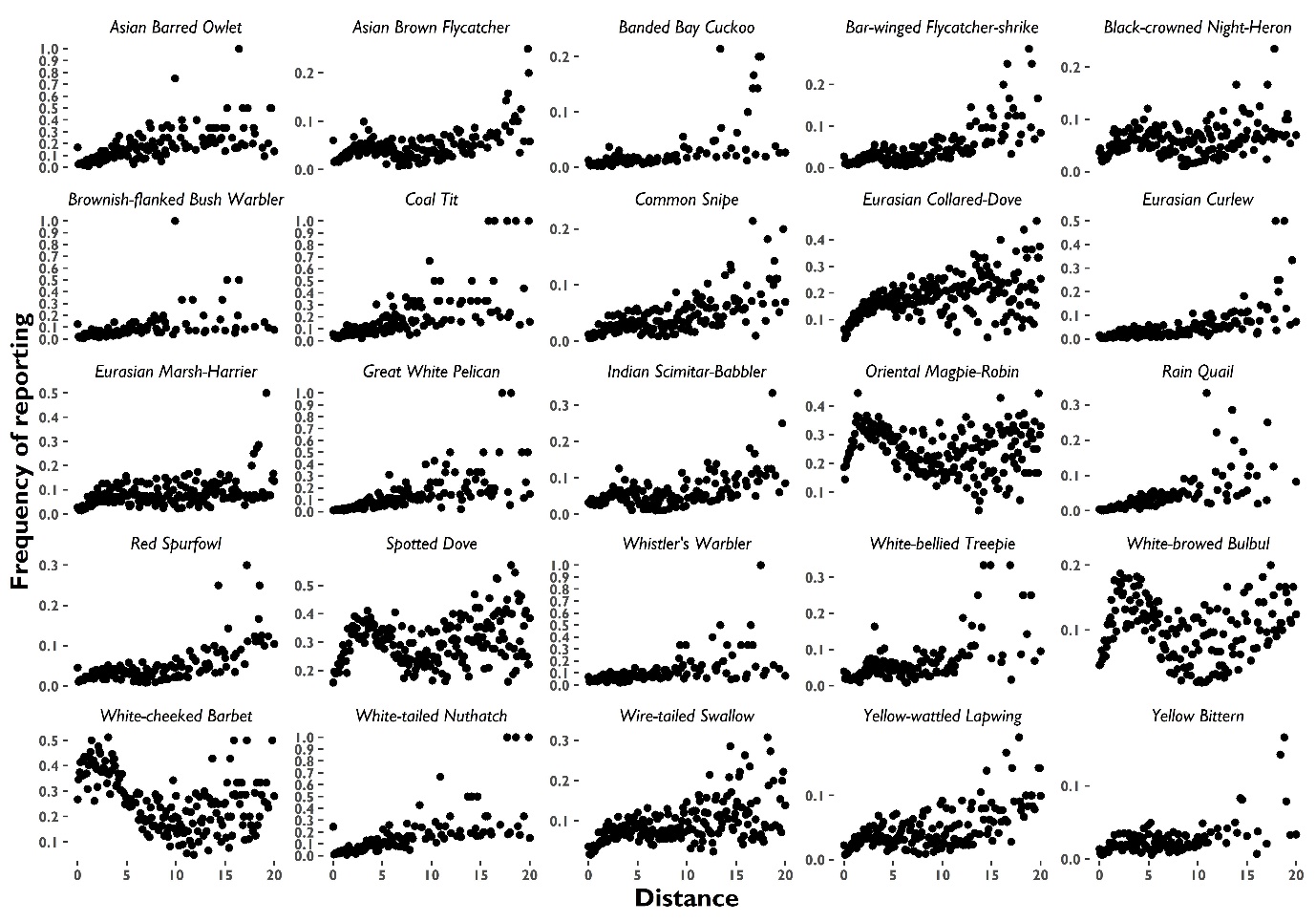

**Fig. S5**: Frequency of reporting of the first 25 of 50 randomly selected species in relation to binned Distance (100 0.2 km bins).

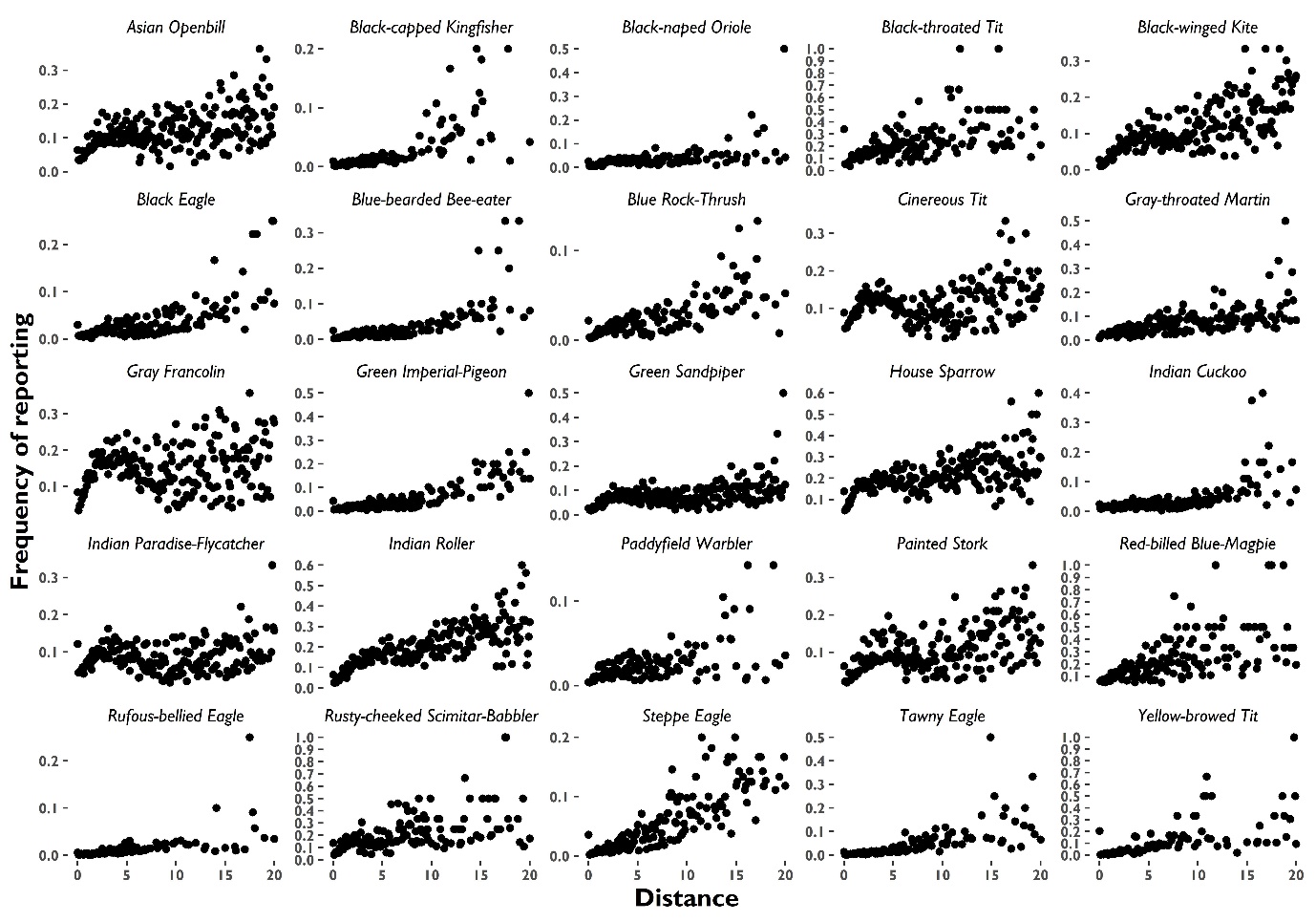

**Fig. S6**: Frequency of reporting of the second 25 of 50 randomly selected species in relation to binned Distance (100 0.2 km bins).

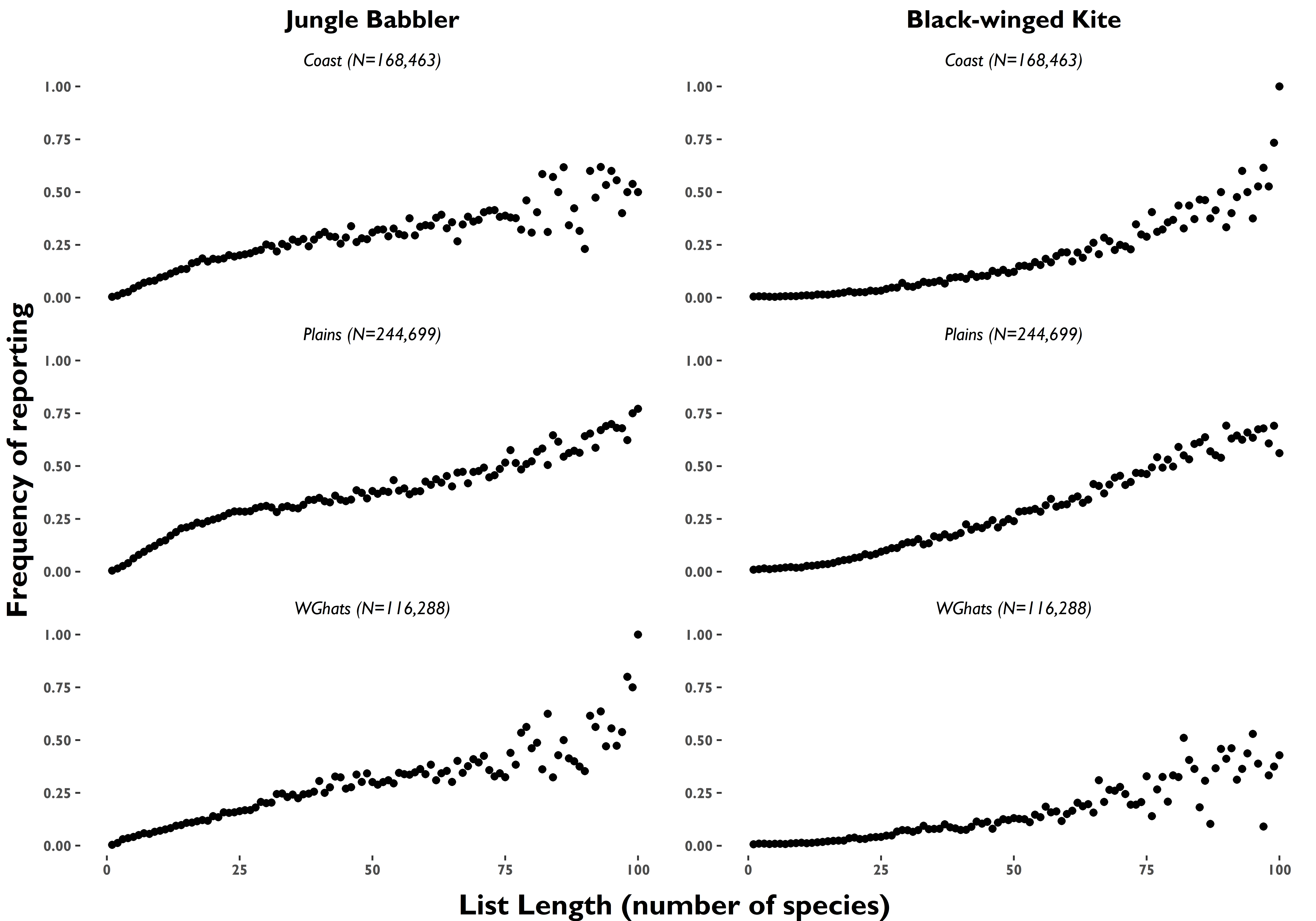

**Fig. S7**: Frequency of reporting of Jungle Babbler and Black-winged Kite in relation to list length (number of species in a checklist up to 100) in three ecological zones: the Coast, Western Ghats, and ‘Rest of peninsula’. Patterns were similar for other widespread species examined separately but not shown here.

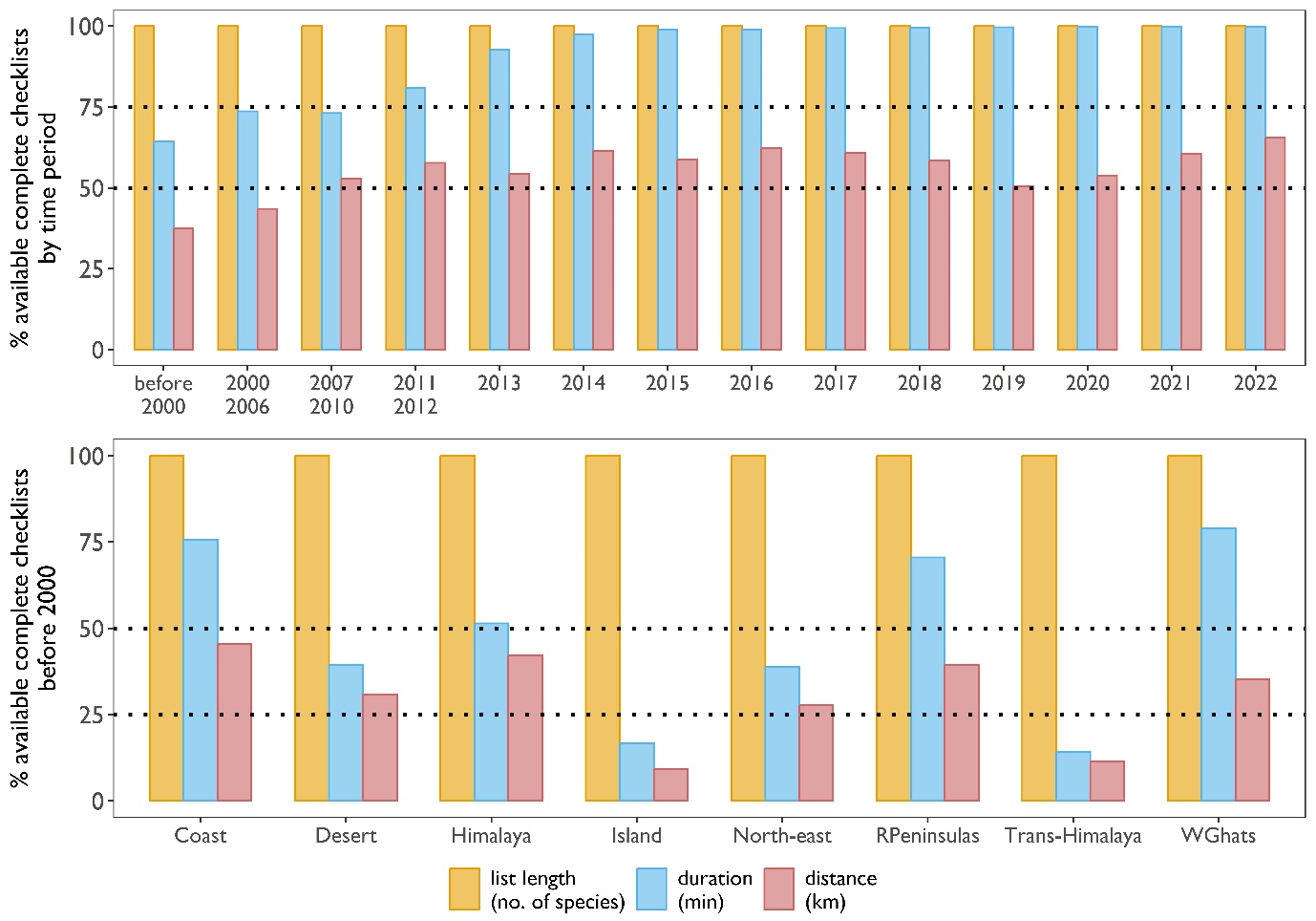

**Fig. S8**: Percentage of data that remains available for analyses when each one of list length, duration and distance travelled are used as covariates, i.e., when excluding data that does not contain each of the three metadata variables. The two panels summarize data availability by year-interval and by ecological region.

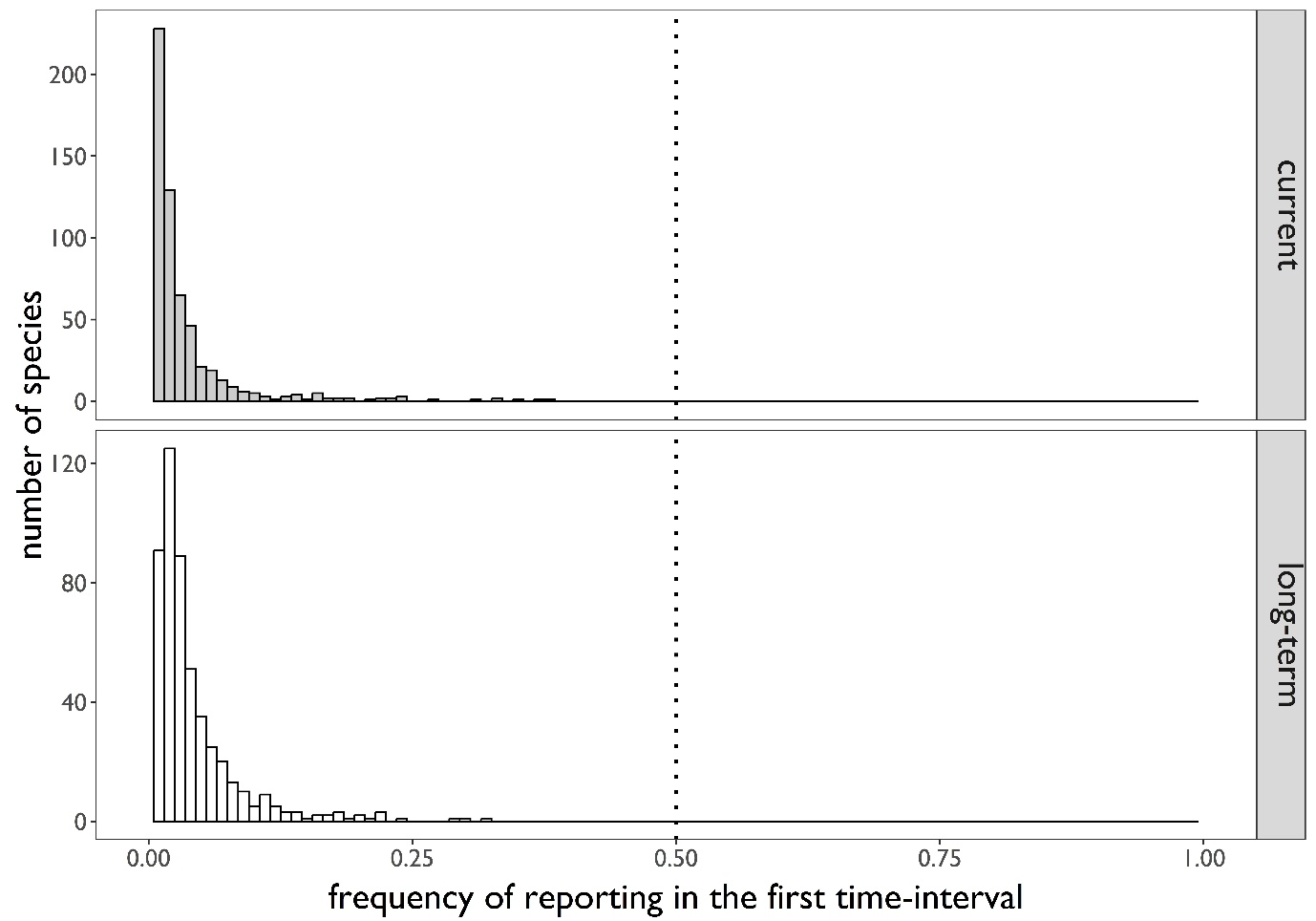

**Fig. S9**: Histograms of species-wise reporting frequencies in the first time-interval (pre-2000 for long-term trends and 2015 for current trend) showing that ratios of the final year with the base time interval can reach at least +100% for all species (given the limit is 1 but all values are less than 0.5).

**Supplementary tables**

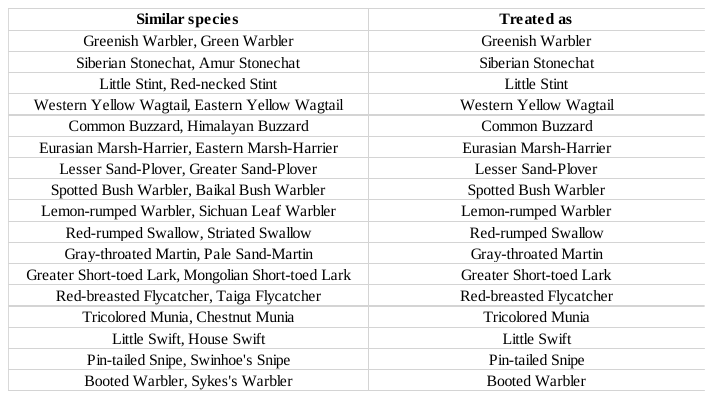

**Table S1**: Similar species that were combined.

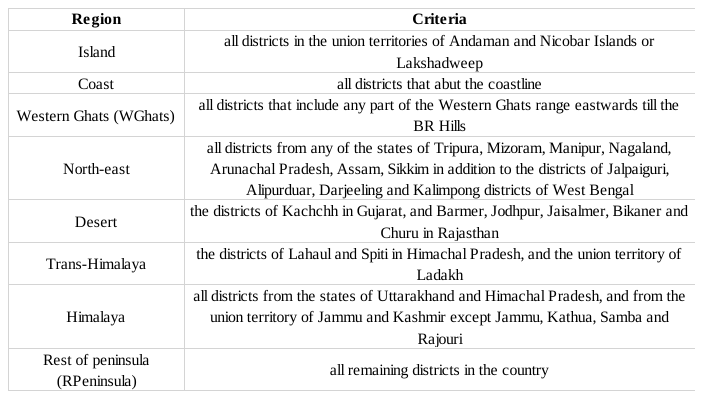

**Table S2**: Criteria to classify Indian administrative districts into any of eight geographical regions.

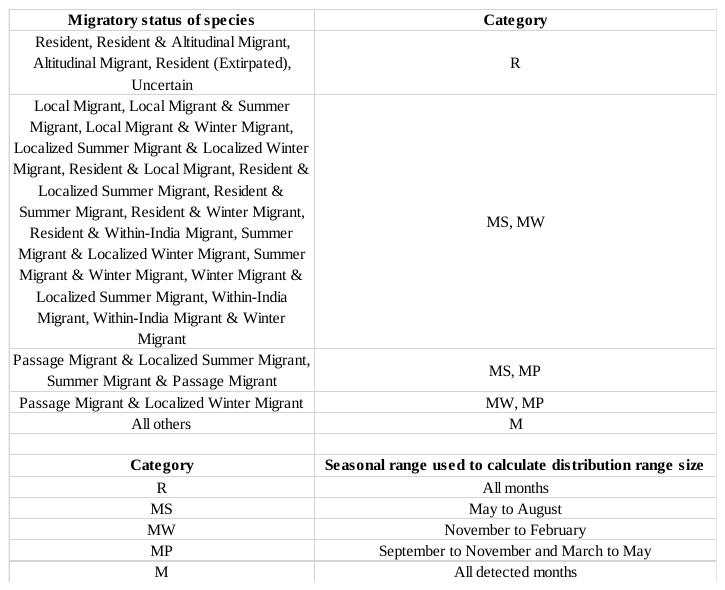

**Table S3**: Categorization of each species based on its migratory status. A species can be in more than one category, which then implies that more than one distribution range size is estimated for that species, each for a different season.

**References**

IUCN. 2022. *The IUCN Red List of Threatened Species*. Version 2022-2. <http://www.iucnredlist.org>. Downloaded on 20 March 2023.

Praveen, J., and R. Jayapal. 2023. Checklist of the birds of India (v7.1). Website: <http://www.indianbirds.in/india/> [Date of publication: 28 February 2023].

SoIB. 2023. *State of India’s Birds, 2023: Range, trends, and conservation status*. Pp. 119. The SoIB Partnership. doi:10.5281/zenodo.11124590.

Wetlands International. 2012. Waterbirds Population Estimates Fifth Edition.

Wildlife (Protection) Act. 1972.
